# Centrosome Loss Triggers a Transcriptional Program to Counter Apoptosis-Induced Oxidative Stress

**DOI:** 10.1101/469056

**Authors:** John S Poulton, Daniel J McKay, Mark Peifer

## Abstract

Centrosomes play a critical role in mitotic spindle assembly through their role in microtubule nucleation and bipolar spindle assembly. Loss of centrosomes can impair the ability of some cells to properly conduct mitotic division, leading to chromosomal instability, cell stress, and aneuploidy. Multiple aspects of the cellular response to mitotic error associated with centrosome loss appears to involve activation of JNK signaling. To further characterize the transcriptional effects of centrosome loss, we compared gene expression profiles of wildtype and acentrosomal cells from Drosophila wing imaginal discs. We found elevation of expression of JNK target genes, which we verified at the protein level. Consistent with this, the upregulated gene set showed significant enrichment for the AP1 consensus DNA binding sequence. We also found significant elevation in expression of genes regulating redox balance. Based on those findings, we examined oxidative stress after centrosome loss, revealing that acentrosomal wing cells have significant increases in reactive oxygen species (ROS). We then performed a candidate genetic screen and found that one of the genes upregulated in acentrosomal cells, G6PD, plays an important role in buffering acentrosomal cells against increased ROS and helps protect those cells from cell death. Our data and other recent studies have revealed a complex network of signaling pathways, transcriptional programs, and cellular processes that epithelial cells use to respond to stressors like mitotic errors to help limit cell damage and maintain normal tissue development.

## Introduction

Proper development requires precise spatial and temporal coordination of cell division to drive tissue growth. During cell division, chromosomes are replicated in S phase, and then segregated equally into two daughter cells during mitosis. The accurate segregation of chromosomes is achieved by the action of the bipolar mitotic spindle (WALCZAK AND HEALD 2008). This microtubule-based structure is essential to generate the physical forces required to move chromosomes to opposite poles, and also has built-in checkpoints that ensure accurate segregation. The assembly of the mitotic spindle is a complex process with multiple layers of regulation to ensure its accuracy (PROSSER AND PELLETIER 2017). Defects in mitotic spindle formation can lead to multipolar spindles or incorrect attachment of microtubules (MTs) to chromosomes, which in turn can lead to segregation errors that cause DNA damage and even whole chromosome mis-segregation (aneuploidy). These types of defects are forms of chromosomal instability (CIN), a hallmark of many cancers that is highly correlated with tumor malignancy (HANAHAN AND WEINBERG 2011; NICHOLSON AND CIMINI 2011).

In most animal cells, the bipolar mitotic spindle arises from the MT nucleating activity of a pair of organelles known as centrosomes, which sit at the two spindle poles (Figure 1A)(WALCZAK AND HEALD 2008; LERIT AND POULTON 2016; PROSSER AND PELLETIER 2017). As the central source of spindle MTs, the orientation of the centrosome pair also determines the geometry of mitotic spindle formation and the axis of division relative to the surrounding tissue. Thus, even within mitosis, centrosomes serve multiple functions related to spindle assembly. Centrosomes also serve a wide range of cellular functions separate from mitotic spindle assembly, including regulation of cilia assembly, cell cycle progression, the DNA damage response, and cell signaling. Given these critical functions ascribed to centrosomes, they were long considered essential components of most animal cells.

**Figure 1.**
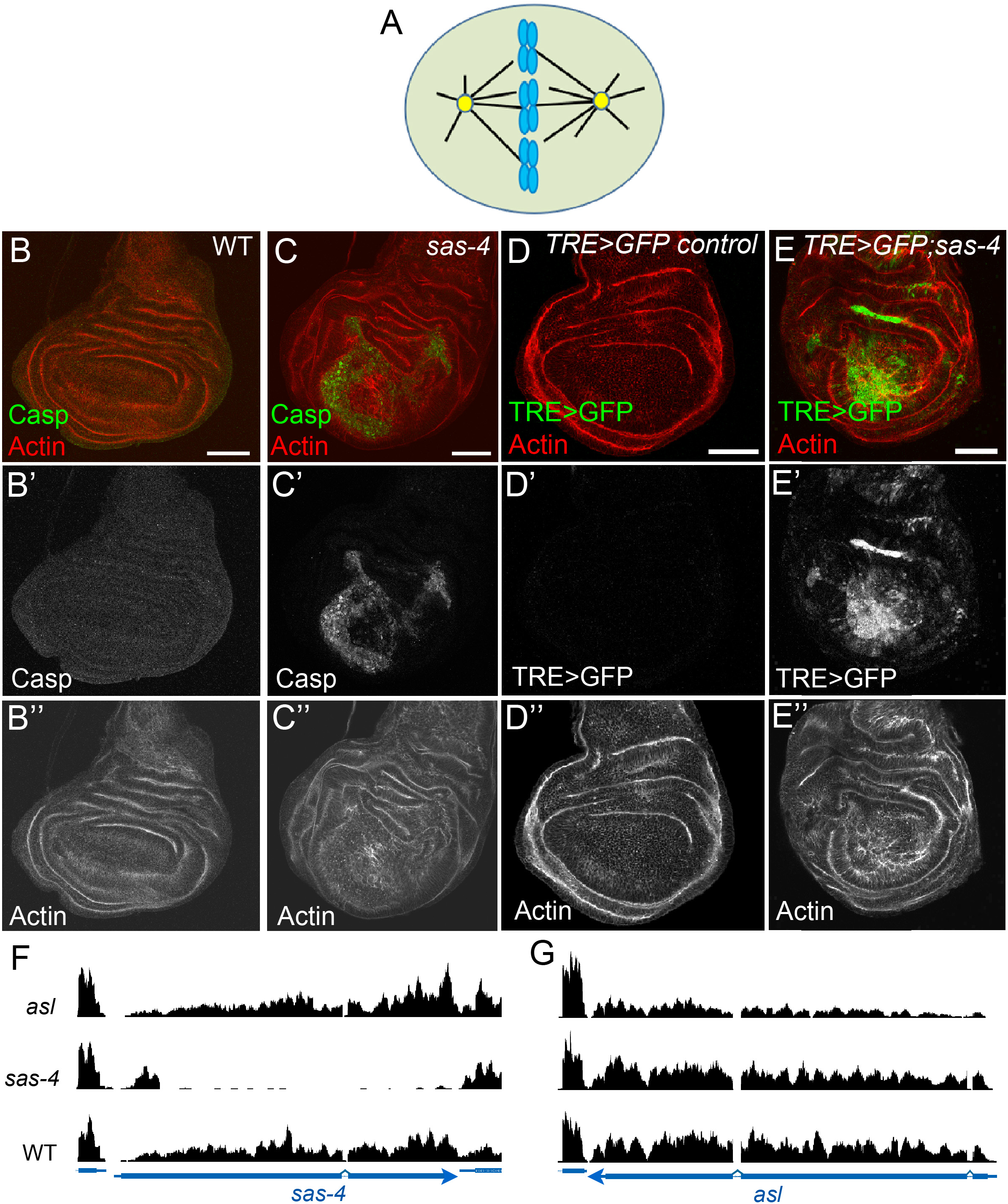
Centrosome loss dramatically increases apoptosis levels and JNK activity. (A) Model of a dividing cell. A pair of centrosomes (yellow) nucleate microtubules (black lines), some of which attach to the chromosomes (blue), to form a bipolar spindle. (B-B”) There is little to no apoptosis in wildtype (WT) 3^rd^ larval instar wing discs, as reported by antibodies to cleaved Caspase 3. (C-C”) In *sas*-*4* mutants, which lack centrosomes, many cells undergo apoptotic cell death. (D-D”) The JNK transcriptional reporter, TRE>GFP, has no detectable expression at this stage in control discs. (E-E”) Many of the acentrosomal cells in *sas*-*4* mutant wing discs have highly elevated JNK activity. All images are maximum intensity projections. Scale bars in all images represent 50μm. Red channel: Actin. Green channel: Cleaved Caspase 3 (Casp) in B,C; TRE>GFP in D,E. (F,G) RNA-Seq reads along the *sas*-*4* locus (F) and *asl* locus (G) in the *asl*, *sas-4*, or WT genotypes.

More recently, however, it has become apparent that cells possess centrosome-independent MT nucleation pathways that assist in spindle assembly (e.g., the Augmin complex and RanGTP pathway)(PROSSER AND PELLETIER 2017). In many cell types, these additional pathways are robust and capable of assembling a bipolar spindle even in the complete absence of centrosomes. A striking example of this occurs in Drosophila where entire animals homozygous mutant for genes essential for centrosome formation or function can develop to adulthood (BASTO *et al*. 2006). We now know this is not unique to flies, because if p53-mediated program cell death is blocked, mice lacking centrosomes can develop to late embryogenesis and then die because the lack of cilia impairs Hedgehog signaling (BAZZI AND ANDERSON 2014).

In Drosophila, detailed examinations of acentrosomal cells in several tissues (e.g., brain and ovarian germline) revealed surprisingly few mitotic errors, indicating the non-centrosomal MT nucleation pathways are adequate for proper spindle assembly and accurate chromosome segregation in those cells (BASTO *et al*. 2006; STEVENS *et al*. 2007; POULTON *et al*. 2017). In contrast to studies in those tissues, we previously found that in the proliferative epithelial cells of the wing imaginal disc, loss of those same centrosomal proteins leads to significant defects in spindle assembly, which increases rates of aneuploidy, DNA damage, and mis-oriented spindles (POULTON *et al*. 2014). Those defects then activate a cell stress pathway, cJun N-terminal Kinase (JNK) signaling, which drives apoptotic cell death (Figure 1B-E). Approximately 15-20% of all cells in acentrosomal wing discs die, suggesting that although alternative MT nucleation pathways help buffer wing disc cells against centrosome loss, they are not as effective in this tissue as they appear to be in other tissues/cell types. Despite the loss of such a substantial fraction of cells, overall wing development remains remarkably normal in most centrosome-deficient animals. Proper mitosis and subsequent wing development in acentrosomal animals are mediated by a number of factors. Correct spindle assembly becomes dependent on MT nucleation by the Augmin complex and RanGTP pathway, and on delay of the cell cycle by the Spindle Assembly Checkpoint (SAC). The cell death that does occur is buffered by compensatory proliferation of neighboring cells to replace dying cells, and delayed development, which presumably allows additional time to correct tissue-level defects caused by massive cell death (POULTON *et al*. 2014). Together, those findings highlighted the remarkable ability of cells and tissues to compensate, not only for loss of key mitotic regulators, such as centrosomes, but also for the wide range of downstream effects of their loss, such as CIN and cell death.

The sensitivity of wing disc cells to mitotic spindle assembly errors due to centrosome loss, as well as their sensitivity to the downstream consequences of spindle assembly errors (i.e., aneuploidy and spindle mis-orientation), make them an excellent model to investigate the cellular response to centrosome loss, mitotic errors, and cell death, as well as the tissue-level and systemic responses to those insults. As our previous data and others demonstrated, one important component of these complex responses to tissue damage induced by a variety of stresses are changes in gene expression, the most well-characterized of which are associated with activation of cell signaling pathways (i.e., JNK, Wnt, Dpp, and JAK-STAT)(RYOO *et al*. 2004; KONDO *et al*. 2006; PASTOR-PAREJA *et al*. 2008; PEREZ-GARIJO *et al*. 2009; DEKANTY *et al*. 2012; POULTON *et al*. 2014). For example, it is now clear that JNK signaling is a central mediator of the response to multiple forms of cell stress or tissue damage (IGAKI 2009). High levels of JNK activity initiates apoptosis in tissues like the wing and eye imaginal discs. To help compensate for the loss of cells due to apoptosis, lower levels of JNK in neighboring cells can help drive proliferation in the surviving cells to help maintain total cell numbers and tissue integrity, which is a central component of the regeneration process (RYOO *et al*. 2004; FAN AND BERGMANN 2008; MARTIN *et al*. 2009; PEREZ-GARIJO *et al*. 2009; FOGARTY *et al*. 2016; BROCK *et al*. 2017; KHAN *et al*. 2017). Several recent studies demonstrate the important transcriptional responses occurring in damaged/stressed cells, much of it mediated directly by JNK signaling. For example, one key pathway that helps drive the compensatory proliferation response is JAK-STAT signaling, whose activating ligands (the Unpaired (Upd) proteins) are themselves transcriptional targets of JNK signaling (PASTOR-PAREJA *et al*. 2008; BUNKER *et al*. 2015; SANTABARBARA-RUIZ *et al*. 2015). Intriguingly, recent studies also uncovered important effects of cell stress and damage on redox balance in imaginal discs, and suggest important roles for reactive oxygen species (ROS) in mediating the activity of the relevant cell signaling pathways to control processes like cell death and compensatory proliferation (KANDA *et al*. 2011; OHSAWA *et al*. 2012; GAURON *et al*. 2013; HUU *et al*. 2015; SANTABARBARA-RUIZ *et al*. 2015; CLEMENTE-RUIZ *et al*. 2016; FOGARTY *et al*. 2016; BROCK *et al*. 2017; KHAN *et al*. 2017; PEREZ *et al*. 2017). Together these studies have begun to elucidate a regulatory network involving complex cross-talk between traditional signaling pathways and ROS that helps correct for cellular damage and maintain tissue homeostasis.

We sought to define the transcriptional response to centrosome loss. To do so, we performed transcriptome analysis on imaginal wing discs from wild type (WT) animals, and from two centrosome-lacking genotypes. Differential gene expression analysis identified hundreds of genes that are significantly up or down-regulated in both acentrosomal mutants relative to WT. One key finding from the transcriptional data, and our subsequent functional genetics experiments, is that centrosome loss induces significant oxidative stress—many of the genes upregulated in acentrosomal cells contribute to redox regulation. We then performed a reverse genetic screen in the genetically sensitized background of acentrosomal wing disc cells and identified a novel genetic interaction between *sas*-*4*, which encodes a core centrosomal protein, and the gene encoding Glucose-6-phosphate dehydrogenase *(g6pd)*, the rate limiting enzyme in the pentose phosphate pathway and key generator of the antioxidant reduced Glutathione (STANTON 2012). We went on to characterize the cellular defects underlying this interaction, and found that G6PD upregulation is an important counter-balance to increased Reactive Oxygen Species (ROS) induced by mitotic errors. Together, the current study reveals new consequences of centrosome loss (i.e., oxidative stress/redox imbalance), as well as yet another way in which acentrosomal cells buffer themselves against the deleterious effects of centrosome loss (i.e., upregulation of antioxidant promoting genes to limit ROS levels).

## Results

### Defining the transcriptional response to centrosome loss

To investigate the effects of centrosome loss on the transcriptional program of a developing tissue, we performed RNA-Seq on *Drosophila* wing imaginal discs from late 3^rd^ instar larvae of three genotypes: *yellow white* (*y w*; our wildtype (WT) control), or animals homozygous mutant for null alleles of one of two different proteins required for centriole duplication: *sas*-*4^s2214^* or *asl^mecD^*. Previous studies demonstrate that these alleles lead to complete or near-complete loss of centrosomes by 3^rd^ larval instar (BASTO *et al*. 2006; BLACHON *et al*. 2008; POULTON *et al*. 2014). We performed RNA-Seq on three biological replicates for each genotype. We first analyzed how well the replicates within a genotype correlated with one another, finding extremely high concordance among replicates for each genotype (Figure 2A; Pearson correlation coefficient=0.99 for replicates within each genotype). We then examined the expression of the *sas*-*4* and *asl* loci in their respective mutant backgrounds. In the *sas*-*4* mutant there was almost complete loss of *sas*-*4* RNA transcripts (Figure 1F). It is worth noting that the *sas*-*4^s2214^* mutant did possess transcripts of ~200bp arising from the 5’ end of the first exon. The *sas*-*4^s2214^* allele is a P-element insertion in the first exon, previously mapped to chromosomal location 3R:2977450, which is precisely where the abrupt end of transcripts was observed in the *sas*-*4* mutant. The *asl^mecD^* mutation is a point mutant (C1718T), leading to a premature stop codon at Q483. Consistent with a point mutant, full length transcripts were present in the *asl^mecD^* mutant background, though overall levels were slightly reduced compared to controls (Figure 1G), which may reflect Nonsense Mediated Decay.

**Figure 2.**
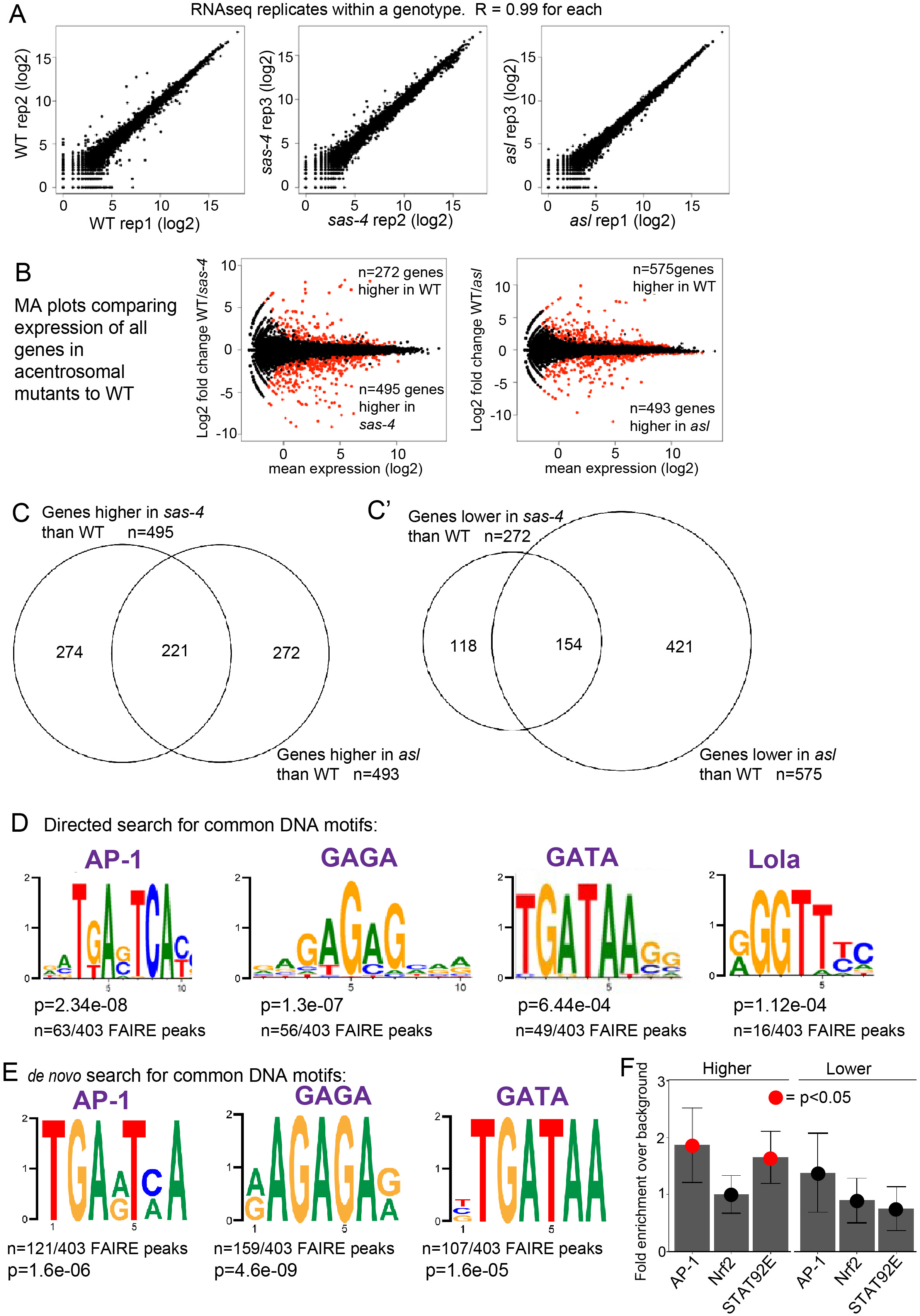
Analysis of RNA-Seq data reveals highly consistent gene up- and down-regulation and enrichment of genes near AP-1, GAGA, GATA, Lola, and STAT92E transcription factor binding sites. (A) Plots comparing RNA-Seq replicates within a genotype demonstrate extremely high concordance. Pearson’s R correlation values are shown. (B) MA plots reveal genes significantly up (red dots below midline) or downregulated (red dots above midline) in comparisons of *sas*-*4* to WT (left) or *asl* to WT (right). (C,C’) Venn diagrams demonstrating the highly significant overlap of genes found to upregulated (C) or downregulated (C’) in both *sas*-*4* and *asl*, relative to WT. (D) Transcription factor DNA-binding motifs significantly enriched in open chromatin sites near to genes upregulated in both *sas*-*4* and *asl*. p-values (rank sum test) and fraction of open chromatin peaks containing a designated motif are shown. (E) A *de novo* motif discovery of open chromatin peaks also uncovered enrichment of AP-1, GAGA, and GATA binding sites. (F) Directed analysis of open chromatin regions near the upregulated and downregulated genes common to *sas*-*4* and *asl* for enrichment of binding sites for AP-1, Nrf2, and STAT92E. Data are plotted as the enrichment over genomic background for each motif. Error bars represent 95% confidence intervals. Red dots indicate a p-value (Z-test) less than 0.05. AP-1 was included as a control for this approach since we knew from our library-based directed search and the *de novo* search that AP-1 binding sites should be significantly enriched in our upregulated set of genes. We did not detect significant enrichment for Nrf2 sites using this approach. However, we did detect significant enrichment of the STAT92E-binding motif in our upregulated genes (this motif was not in the library for our directed search, represented in panel D).

To define changes in gene expression associated with centrosome loss, we compared RNA-Seq data from each of the acentrosomal mutants to the WT control RNA-Seq data. Plots of differential gene expression revealed many up and down-regulated genes for each pairwise genotype comparison (Figure 2A; complete lists of gene expression data are found in Suppl. Table 1). To identify the genes most significantly up or down-regulated in each mutant relative to WT, we filtered the comprehensive list of genes to include only those genes that met a False Discovery Rate (FDR) of p<0.001, as well as a minimum expression threshold (FPKM ≥10) for at least one genotype. In the comparison of *sas*-*4* mutant wing discs to controls, use of these filters revealed 495 genes significantly upregulated and 272 genes downregulated in *sas*-*4* discs (Suppl. Table 2). In the comparison of *asl* mutant wing discs to controls, we found 493 upregulated genes and 575 downregulated (Suppl. Table 3).

Mutations in *sas*-*4* and *asl* result in centrosome loss through different mechanisms—Sas-4 is directly involved in centriole assembly (KOHLMAIER *et al*. 2009; SCHMIDT *et al*. 2009), whereas Asl regulates daughter centriole duplication licensing (BLACHON *et al*. 2008; NOVAK *et al*. 2014). Therefore, changes in gene expression unique to one mutant genotype might reflect the transcriptional response to centrosome-independent functions for that particular protein. To identify the common response to centrosome loss, we cross-referenced the lists of differentially-expressed genes to identify sets of genes that were significantly up or down-regulated in <u>both</u> acentrosomal mutants relative to WT. The two lists exhibited highly significant overlap (Figure 2C,C’): 221 genes were significantly upregulated (Suppl. Table 4) and 154 genes were significantly downregulated (Suppl. Table 5) in <u>both</u> *sas*-*4* and *asl* wing discs, relative to control wing discs. This was much higher than expected by chance (hypergeometric mean test: p<4.26^e−201^ for upregulated genes; p<1.54^e−91^ for downregulated genes). This conservative approach should exclude any unforeseen changes in gene expression associated with a particular mutant or mutant background, thus isolating only genes specifically affected by loss of centrosomes. Of course, for some genes, lack of concordance may simply result from experimental variability. Similarly, changes in gene expression for a particular gene might have reached the significance threshold in one genotype, but been just below that threshold in the other, thus excluding it from the shared list of significantly up or downregulated genes common to both.

Several categories of genes, as defined by GO-Term analysis via DAVID (HUANGDA *et al*. 2009), were notable in the list of jointly upregulated genes. These included genes involved in oxidation-reduction pathways, including the antioxidant and detoxifying glutathione transferase pathway, as well as genes involved in the innate immune response (Tables 1-3). Manual inspection also revealed a number of genes involved in or known to be targets of the JNK pathway (Table 4), consistent with our previous work (POULTON *et al*. 2014).

**Table 1.** GO Terms enriched in genes higher in both *sas*-*4* and *asl*. Gene Ontology (GO) Term analysis of genes with significantly increased expression in both *sas*-*4* and *asl* mutant wing discs, relative to WT, suggests upregulation of several biological pathways. Notable among them are indicators of oxidative stress. Unadjusted p-values are shown.

| **<u>Biological Process</u>** | **<u>p-value</u>** |
| --- | --- |
| Glutathione Metabolic Process | 3.90E-04 |
| Imaginal Disc-derived Male Genitalia Morphogenesis | 1.20E-03 |
| Cell Adhesion | 1.60E-03 |
| Immune Response | 5.80E-03 |
| Oxidation-reduction Process | 9.20E-03 |
| **<u>Molecular Function</u>** | **<u>p-value</u>** |
| Glutathione Peroxidase Activity | 8.45E-05 |
| Signaling Pattern Recognition Receptor Activity | 3.30E-03 |

### The transcriptional response to centrosome loss does not broadly elevate core centrosomal proteins or proteins involved in parallel pathways

Centrosomes are multi-protein organelles. We thus had initially hypothesized that cells might sense the mitotic challenge in centrosome-deficient cells by upregulating the genes encoding centrosomal proteins. However, no known centrosomal components were on the list of genes significantly upregulated in both *asl* mutants and *sas*-*4* mutants. Centrosome loss in wing imaginal discs is buffered by the mitotic delay induced by the SAC and by the microtubule nucleation that mediates non-centrosomal spindle assembly. Thus another potential transcriptional response might be upregulation of the components of the SAC, Augmin complex, or Ran pathway, which partially compensate for centrosome loss in wing discs as well as in the early embryo (HAYWARD *et al*. 2014; POULTON *et al*. 2014). Only two genes with microtubule or SAC connections were on the list of genes significantly upregulated by loss of both Asl and Sas-4: Tubulin binding cofactor A (CG1890)(VOELZMANN *et al*. 2016), and Spindly, a protein essential for silencing the SAC via dynein recruitment to the kinetochore (GRIFFIS *et al*. 2007). However, when we scanned the lists of genes upregulated by knockdown of *sas4* or *asl* alone, a few additional genes emerged: *rcd2*, identified in an RNAi screen for centrosome function (DOBBELAERE *et al*. 2008), and *CP309*, encoding the centrosomal protein Pericentrin-like protein (PLP)(MENNELLA *et al*. 2012; LERIT *et al*. 2015; RICHENS *et al*. 2015), were upregulated in *sas*-*4* mutants, while *mad2*, a key component of the spindle assembly checkpoint (MUSACCHIO 2015), *ran*, which has dual roles in nuclear import and in non-centrosomal microtubule nucleation (CLARKE AND ZHANG 2008), and *cct5*, involved in centrosome-independent spindle assembly (MOUTINHO-PEREIRA *et al*. 2013), were upregulated in *asl* mutants. Thus coordinated transcriptional upregulation of the compensatory pathways does not appear to be a prominent response to loss of centrosomes, but it may play a minor role.

Centrosome loss in the wing imaginal disc disrupts mitotic spindle assembly, leading to chromosome mis-segregation and DNA damage (POULTON *et al*. 2014). The DNA damage response (DDR) is a complex, multi-tiered process wherein damage to DNA initiates repair pathways to correct the lesions, or, if the damage is too severe, triggers programmed cell death (BORGES *et al*. 2008). One aspect of the DDR is upregulation of genes involved in DNA damage detection or repair (CHRISTMANN AND KAINA 2013). Interestingly, we did not detect significant up or downregulation of known DDR genes in our analysis. Among the many possible explanations for this are that the extent of DNA damage is not sufficient to detect by analysis of the entire tissue (DNA damage was only detected in a subset of cells in acentrosomal wing discs, possibly due to rapid elimination of damaged cells from the wing epithelium)(POULTON *et al*. 2014), or that changes in gene expression are not a primary component of the DDR in this tissue.

### Validating differential gene expression associated with centrosome loss reveals significant upregulation of genes regulated by the JNK pathway

In wing discs lacking centrosomes (i.e., *sas*-*4* mutant), there are significant defects in efficient spindle assembly, accurate chromosome segregation, and proper spindle orientation, leading to increased apoptosis (Figure 1B,C)(POULTON *et al*. 2014). These mitotic defects appear to drive apoptosis of affected cells by activation of JNK signaling, since blocking JNK signaling prevented apoptosis in acentrosomal wing discs (POULTON *et al*. 2014). JNK signaling regulates gene expression, at least in part through the key transcription factor AP-1, a heterodimer of Jun (Jun-related antigen (Jra) in flies) and Fos. Consistent with this, we previously found that a JNK signaling transcriptional reporter (TRE>GFP; expresses GFP under the control of a promoter containing Jun binding sites) is activated by centrosome loss (i.e., *sas*-*4* RNAi), both in cells undergoing apoptosis and also in other cells in the disc (Figure 1D,E)(POULTON *et al*. 2014). These data suggested that we would see elevated expression of JNK target genes in centrosome deficient discs, both at the transcript and protein levels.

We thus tested this hypothesis, focusing on the *sas*-*4* mutant because *sas*-*4* loss appeared to elicit a stronger transcriptional response than *asl* for many of the genes present on the shared list of differentially expressed genes. Many positively regulated transcriptional targets of JNK signaling have been identified in *Drosophila*. Consistently, mRNA levels of many of these, including *Jra* itself, *puckered* (*puc*), a feedback negative regulator of the JNK pathway, *Insulin*-*like peptide 8* (*Ilp8*), *Reaper* (*rpr*), *Matrix Metalloproteinase1* (*MMP1*), were elevated in both the *sas*-*4* and *asl* mutant backgrounds, relative to WT controls (Figure 3A,B,E,F,L,M; Suppl Table 4).

**Figure 3.**
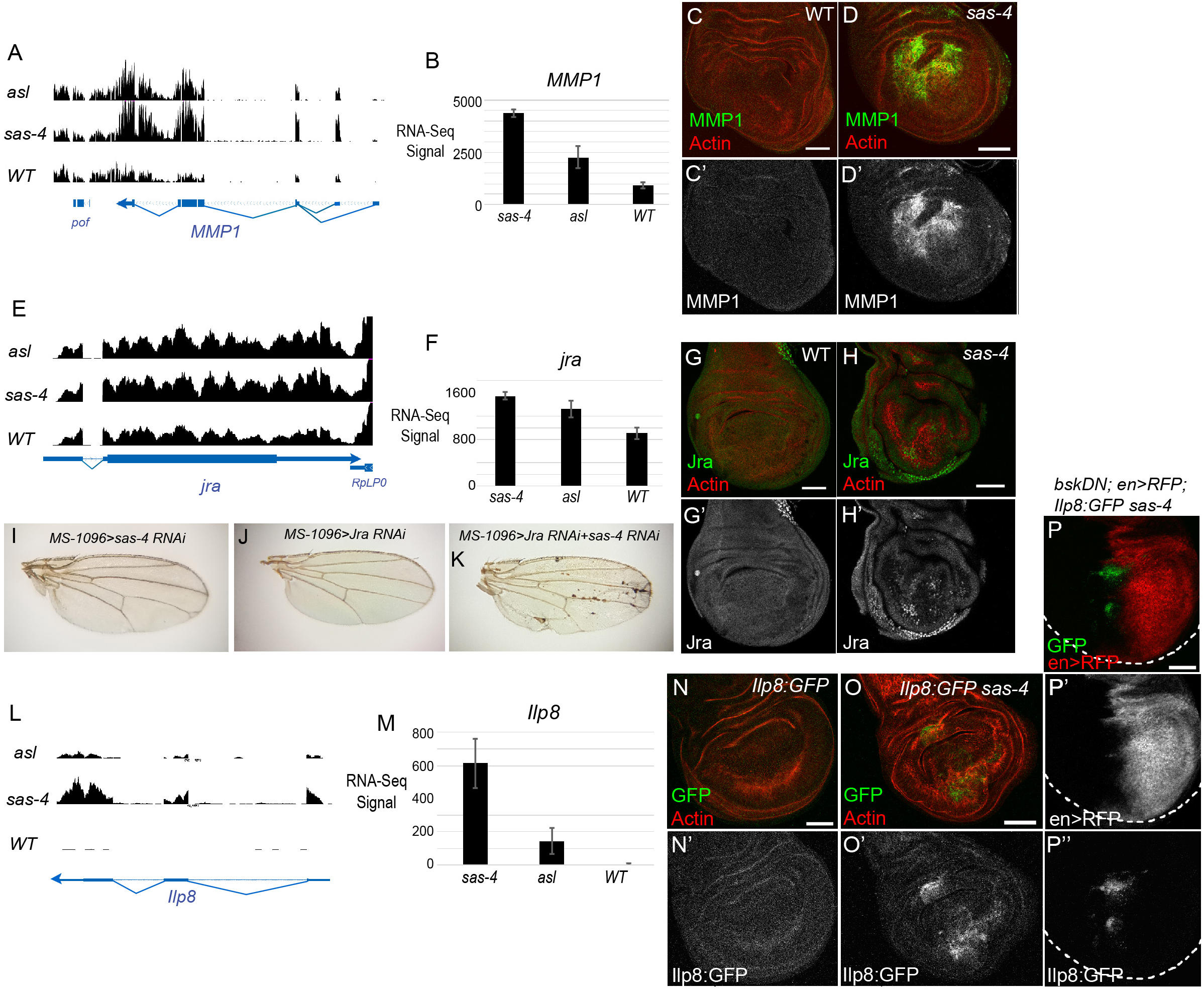
Centrosome loss leads to upregulation of expression of JNK target genes. (A,E) RNA-Seq plot profiles of two known JNK target genes, *MMP1* (A) and *Jra* (E), for *asl*, *sas*-*4* and WT genotypes. Transcription direction indicated by arrowheads. (B,F) RNA-Seq mean signal (+/- s.d.) of the three biological replicates for *MMP1* (B) and *Jra* (F). (C) MMP1 protein, as visualized by antibody, has minimal expression in WT wing discs. (D) MMP1 protein levels are dramatically increased in acentrosomal *sas*-*4* wing discs. (G) Jra is weakly expressed in control discs, though there is significant expression in the peripodial cells (not shown in this single slice image). (H) Jra protein is increased in *sas*-*4* discs. (I-K) Representative adult wings from the indicated genotypes. Neither knockdown of *sas*-*4* alone (I) nor *jra* alone (J) perturbs wing development. However, knockdown of both *sas*-*4* and *jra* together produces necrotic spots in the adult wing. (L) RNA-Seq plot profiles of the *ilp8* locus for *asl*, *sas*-*4* and WT genotypes. (M) Mean RNA-Seq signal for *ilp8* in the three genotypes. (N) *ilp8* expression, as assessed using a protein trap line expressing GFP tagged Ilp8 under control of the *ilp8* promoter, is low in controls discs. (O) *ilp8* is upregulated in *sas*-*4* discs. (P) The upregulation of *ilp8* associated with centrosome loss is JNK-dependent because misexpression of BskDN in the posterior portion of *sas*-*4* homozygous mutant wing discs inhibits Ilp8:GFP upregulation. BskDN is driven by en>RFP (red in P; grayscale in P’). White dashed line marks the outer edge of the wing disc. Scale bars are 50μm. Images are max projections, except in G,H where single slices were used to limit the Jra signal from the peripodial cells.

We next examined whether the changes in RNA transcript levels observed in RNA-Seq analysis led to changes in protein levels of JNK transcriptional-targets, by using antibodies to MMP1 and Jra. Control WT wing discs express little to no MMP1or Jra protein (Figure 3C,G; with the exception of peripodial cells, which express moderate levels of Jra). Consistent with the RNA-Seq data, we found that *sas*-*4* mutant discs had noticeable increases in both proteins (Figure 3D,H). Because Jra is an essential component of the JNK signaling pathway, we tested the importance of Jra expression in control and acentrosomal cells. While knocking down Jra or Sas-4 alone did not perturb wing development, knocking down both led to significant morphological defects (Figure 3I-K). This is consistent with our previous data demonstrating a role for JNK itself in maintaining tissue homeostasis in acentrosomal wing discs (POULTON *et al*. 2014). This interaction likely occurs through JNK’s positive roles in apoptosis and/or compensatory proliferation stemming from centrosome loss.

Another interesting hit from our RNA-Seq screen was *Insulin*-*like peptide 8* (*Ilp8*)(Figure 3L,M; Suppl Table 4). Ilp8 mediates delays in developmental timing caused by abnormal tissue growth during larval stages (COLOMBANI *et al*. 2012; GARELLI *et al*. 2012). We found that larvae mutant for centrosomal proteins such as Sas-4 exhibit a significant delay in larval development, taking approximately 24hrs longer than controls to enter pupation (POULTON *et al*. 2014). We thus examined whether the increase in Ilp8 transcripts in centrosome-deficient animals leads to increased Ilp8 protein expression, using Ilp8:GFP, a GFP protein trap of the endogenous *Ilp8* locus (GARELLI *et al*. 2012). In control WT animals, there is minimal expression of Ilp8 in 3^rd^ instar wing imaginal discs (Figure 3N). However, in *Ilp8:GFP sas*-*4* animals, we noted a significant increase in Ilp8:GFP expression (Figure 3O). Ilp8 upregulation in response to imaginal disc growth defects induced by knockdown of endocytic or ribosomal proteins (i.e., Avl or Rpl7) requires JNK signaling (COLOMBANI *et al*. 2012). We thus tested whether JNK signaling mediated the upregulation of Ilp8 after centrosome loss. Indeed, when we used the Gal4-UAS system to ectopically express a dominant negative form of the fly homologue of JNK (BasketDN; BskDN) in the posterior portion of *sas*-*4* homozygous mutant wing discs, this led to a clear reduction in Ilp8:GFP in the region of the disc where JNK was inhibited (Figure 3P). The increased Ilp8 expression in *sas*-*4* mutants, along with the known developmental delay and JNK activation experienced by these animals, suggests Ilp8 upregulation via JNK is likely an important mediator of prolonged development in acentrosomal animals.

These data demonstrate that, in the wing imaginal disc, centrosome loss leads to increased JNK activity with concomitant changes in expression of JNK target genes, and also validate the accuracy of our RNA-Seq data. It will be interesting to determine if some of the other genes differentially regulated by centrosome loss, as identified in our RNA-Seq data, are previously unknown JNK signaling targets—a growing number of transcriptomic studies from Drosophila models with active JNK signaling may provide valuable data for cross-referencing (ROUSSET *et al*. 2010; BUNKER *et al*. 2015; CLEMENTE-RUIZ *et al*. 2016; KHAN *et al*. 2017).

To examine this possibility in our own data, we performed transcription factor binding motif analysis of the shared genes upregulated in acentrosomal cells. We first looked for enrichment of known transcription factor binding motifs in open chromatin sites of 3^rd^ instar wing imaginal discs within 2kb of the 221 genes upregulated in both mutant backgrounds (=403 FAIRE peaks)(UYEHARA *et al*. 2017). Remarkably, a consensus sequence significantly matching the AP-1 binding site was found in 63 of the 403 FAIRE peaks (Figure 2D; p=2.34^e-8^). This suggests that there may be many additional genes directly upregulated by JNK signaling in centrosome-deficient cells. To further test this possibility, we performed *de novo* motif discovery in those open chromatin regions within 2kb of the upregulated genes. This analysis revealed the presence of an AP-1 binding site in 121/403 FAIRE peaks (Figure 2E; p=1.6^e-6^). Intriguingly, these analyses also revealed additional motifs unrelated to JNK signaling, including GAGA, GATA, and Lola binding sites (Figure 2D,E). Notably, GATA proteins, including the primary fly GATA protein Serpent, have recently been found to be regulated by ROS levels (GAO *et al*. 2014; INDO *et al*. 2017), and help promote the innate immune response (SENGER *et al*. 2006)—as noted above, centrosome loss increases expression of genes involved in oxidative stress and innate immune responses (Table 1). It will be interesting in the future to test possible roles for these transcription factors in the response to centrosome loss. It is also worth noting that neither *de novo* nor motif-enrichment analyses applied to genes downregulated in acentrosomal cells revealed any significant support for particular transcription factor binding sites near to those genes.

Based on the upregulation of the JAK-STAT ligands Upd2 and Upd3 in acentrosomal cells (Suppl. Table 4), it was curious that the STAT92E binding motif was not found in our *de novo* search (Figure 2E; the STAT92E consensus sequence was absent from the library of motifs used in our directed search). We therefore conducted a third analysis of potential transcription factor binding sites in the open chromatin around the genes up or downregulated in *sas*-*4* and *asl*, this time specifically looking for enrichment of sequences aligning to the STAT92E binding motif. In this analysis, we used the consensus AP-1 binding motif as a positive control since it was found to be significantly enriched in the upregulated genes based on both of our other motif search approaches (Figure 2D,E). This analysis revealed significant enrichment of STAT92E binding sites in the open chromatin regions of genes upregulated by centrosome loss (Figure 2F). In contrast, there was no significant enrichment of STAT92E binding sites in genes downregulated by centrosome loss. To more directly determine whether *sas*-*4* knockdown leads to upregulation of JAK-STAT activity in acentrosomal cells, we examined expression of the JAK-STAT transcriptional reporter 10xSTAT:GFP in *sas*-*4* mutant wing discs. While, we did not detect obvious changes in JAK-STAT activity in *sas*-*4* mutants, when we blocked apoptosis with p35, we did find increased JAK-STAT activity (Suppl Figure 1), suggesting that centrosome loss leads to increased JAK-STAT activation, likely through JNK-induced upregulation of Upd ligands. Intriguingly, the strongest upregulation was in neighboring wildtype cells (Suppl Figure 1C, arrows), which may reflect JAK-STAT’s involvement in the compensatory proliferation response.

### Centrosome loss leads to oxidative stress

One of the most striking features of our RNA-Seq data was increased expression of a number of genes associated with the response to oxidative stress (Table 2). These ranged from signaling proteins like TNF-associated Factor 4 (TRAF4), which acts upstream of the JNK pathway to regulate the oxidative stress response (TANG *et al*. 2013), to enzymes like WW domain containing oxidoreductase (WWOX), which regulates ROS and TNF-induced cell death (O’KEEFE *et al*. 2015), or CG3714, a Nicotinate phosphoribosyltransferase family member that is essential for the increase in cellular NAD levels to prevent oxidative stress (HARA *et al*. 2007). Among these, multiple Glutathione S-transferase (GST) genes were upregulated in both *sas*-*4* and *asl* mutants, and *sas*-*4* loss led to upregulation of three additional GSTD genes (Figure 4A). These enzymes mitigate oxidative stress by conjugating glutathione to toxic electrophilic substrates, reducing their reactivity and increasing their solubility, thus facilitating their excretion from cells and tissues (CHATTERJEE AND GUPTA 2018).

**Figure 4.**
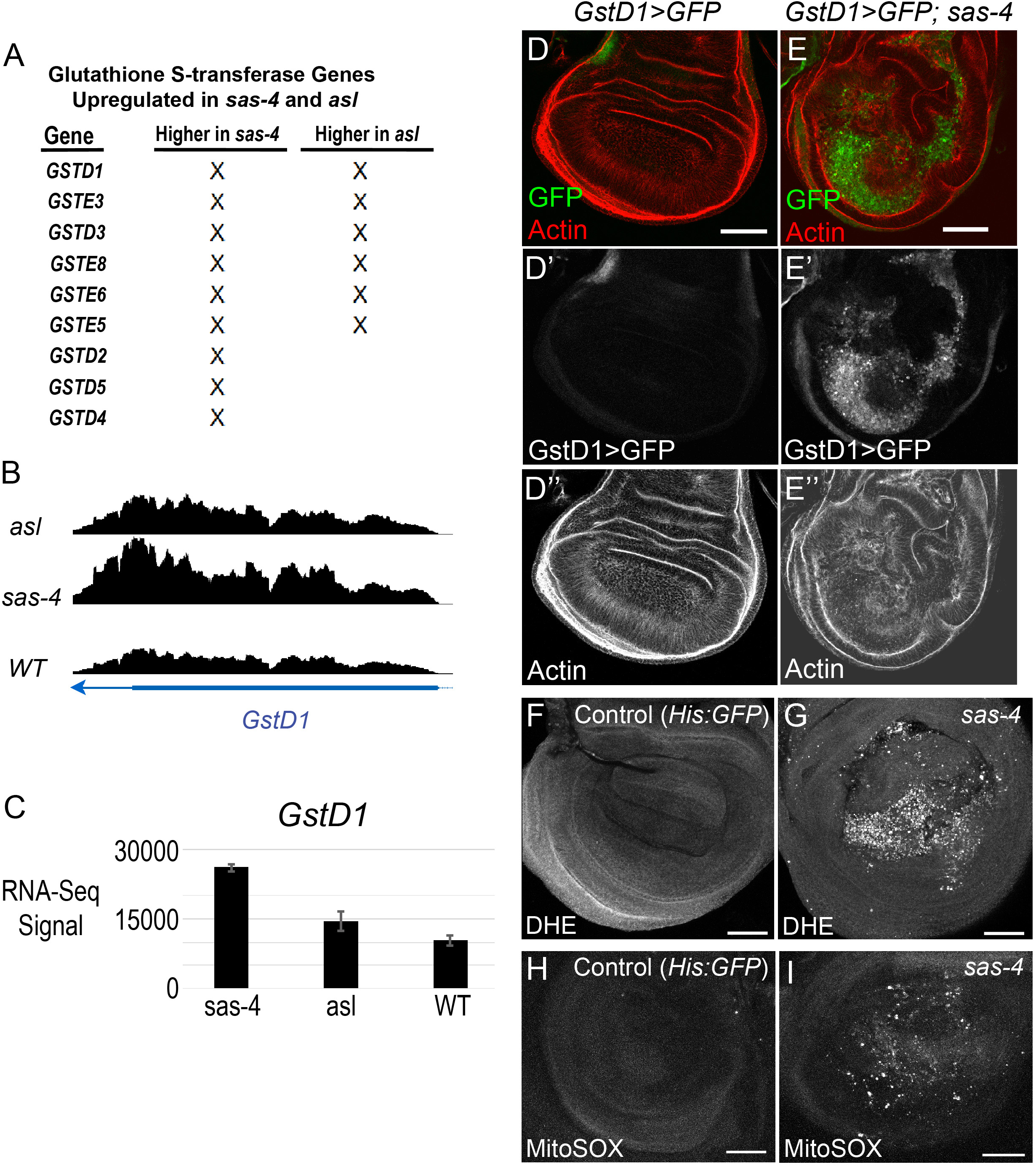
Centrosome loss induces oxidative stress and upregulation of GST genes. (A) List of *GST* genes significantly upregulated in *sas*-*4* and/or *asl* mutants. (B) RNA-Seq plot profiles of the *GstD1* locus in the three genotypes. Transcription direction indicated by arrowhead. (C) Mean RNA-Seq signal for *GstD1*. (D) Control WT wing discs have minimal expression of *GstD1*, as indicated by the reporter GstD1>GFP. (E) *sas*-*4* wing discs have elevated levels of GstD1>GFP. (F) In WT wing discs, ROS levels are essentially undetectable. His:GFP flies were used as WT in this experiment, and were mixed with *sas*-*4* mutant discs to provide an “in tube” control. (G) *sas*-*4* homozygous mutant wing discs showed strongly elevated ROS levels. (H) WT wing discs do not stain for MitoSOX, a marker of mitochondrially-derived super-oxide. (I) MitoSOX staining is elevated in *sas*-*4* mutant discs. Scale bars are 50μm.

**Table 2.** Genes with known or putative roles in the oxidative stress response. A list of oxidative stress response genes that were significantly upregulated in both *sas*-*4* and *asl* mutants relative to WT.

| <u>Gene</u> | <u>Protein Name/Function</u> | <u>Reference</u> |
| --- | --- | --- |
| Zw (G6PD) | Zwischenferment. Glucose-6-phosphate dehydrogenase. Rate-limiting enzyme in the pentose phosphate pathway. Reaction generates NADPH, which is used by Glutathione reductase to produce reduced Glutathione (GSH), a potent antioxidant. A Nrf2 target. | Loboda et al, 2016 |
| Glutathione synthase | Second enzyme in synthesis of Glutathione, the key non-enzymatic antioxidant. | Lu, 2013 |
| Traf4 | TNF-receptor-associated factor 4. Traf2 regulates oxidative stress with Atg9 through JNK. | Tang et al, 2013 |
| Wwox | WW domain containing oxidoreductase. Regulates ROS and TNF-induced cell death. | O'Keefe et al, 2015 |
| Aox (CG18522 ) | Aldehyde oxidase 1. Potent generator of superoxides. | Kundu et al, 2012 |
| GstE8 | Glutathione-S-transferase E8. Cnc and Paraquat induced. Up in hyperoxia screen. | Greunewald et al, 2009; Misra et al, 2011 |
| GstD3 | Glutathione S transferase D3. Cnc and Paraquat induced. Up in hyperoxia screen. | Greunewald et al, 2009; Misra et al, 2011 |
| Men | Malic oxidoreductase. Malic enzyme knockdown modulates reductive stress. | Xie et al., 2013 |
| ImpL3 | Lactate dehydrogenase. HIF-1 target gene. Up in hyperoxia screen. Inhibition induces oxidative stress. | Greunewald et al, 2009 |
| Naprt (CG3714) | Nicotinate phosphoribosyltransferase. Rate limiting enzyme in NAD<sup>+</sup> synthesis. NAD<sup>+</sup> is a cofactor in redox reactions and helps prevent oxidative stress. | Massudi et al, 2013 |
| AcCoAS | Acetyl Coenzyme A synthase. Mammalian Acss2 promotes acetylation of the stress-responsive Hypoxia Inducible Factor 2α (HIF-2α) subunit by the acetyltransferase/coactivator Creb binding protein (Cbp). | Chen et al, 2017 |
| CG3397 | Putative Aldo/keto reductase. Up after hyperoxia oxidative stress. | Greunewald et al, 2009 |

We thus followed this lead, using GSTD1 as an example. GSTD1 is a known target of KEAP1/Nrf2 signaling, which regulates the response to oxidative stress (SYKIOTIS AND BOHMANN 2008). Experiments in the eye imaginal disc indicate GSTD1 can also be upregulated by JNK signaling (KANDA *et al*. 2011). Our RNA-seq data suggested that GSTD1 is upregulated in acentrosomal cells (Figure 4A-C). To confirm this, we took advantage of a GSTD1>GFP reporter, in which the promoter region of GSTD1 drives GFP expression (SYKIOTIS AND BOHMANN 2008). In WT discs, GSTD>GFP expression is very low, while in *sas*-*4* mutant discs, there is strong upregulation of GFP driven by the GSTD1 promoter (Figure 4D,E). Interestingly, the consensus binding site for Nrf2 did not show up in our *de novo* transcription factor motif analysis (Figure 2E), nor was it found to be significantly enriched in our directed motif search of genes up or downregulated by centrosome loss (Figure 2F). Thus, despite significant upregulation of several oxidative stress response genes, including GSTD1, the number of direct targets of KEAP1/Nrf2 signaling in our upregulated gene set may be rather small.

Increased expression of GSTD genes, and in particular the upregulation of GSTD1>GFP, can reflect the presence of ROS (SYKIOTIS AND BOHMANN 2008). As noted above, Gene Ontology (GO) term analysis of our lists of genes significantly up or down-regulated in acentrosomal cells revealed changes in expression of proteins involved in both redox metabolism and detoxification associated with xenobiotic factors and oxidative stress. This could reflect an increase in reactive oxygen species (ROS) in cells lacking centrosomes. To test this hypothesis, we incubated *sas*-*4* mutant wing discs with the ROS probe dihydroethidium (DHE)(BINDOKAS *et al*. 1996). Strikingly, *sas*-*4* mutant wing discs had a dramatic increase in DHE staining (Figure 4F,G). We also observed increased signal using the MitoSOX probe, indicating at least some ROS production occurs in mitochondria (Figure 4H,I).

Centrosomes regulate several cellular processes that could conceivably affect redox balance (e.g., the DNA damage response)(LERIT AND POULTON 2016). We therefore sought to determine if the increase in ROS was a specific effect of centrosome loss, or a potential consequence of the mitotic errors induced by centrosome loss in the wing disc. Knockdown of key mitotic regulators and resulting mitotic errors were recently reported to lead to increased GSTD>GFP expression (CLEMENTE-RUIZ *et al*. 2016). We therefore used our ROS assays to examine ROS levels following knockdown of two other mitotic regulators, Mud and Bub3. Mud is important for spindle orientation in the wing disc (NAKAJIMA *et al*. 2013), while Bub3 contributes to the Spindle Assembly Checkpoint and the attachment of MTs to kinetochores (LOGARINHO *et al*. 2008). Thus, defects caused by knockdown of these proteins should be independent of centrosome function. As previously reported (DEKANTY *et al*. 2012; MORAIS DA SILVA *et al*. 2013; POULTON *et al*. 2014), knockdown of each of these proteins leads to significant increases in apoptosis (Figure 5A,C). We found that Mud or Bub3 knockdown also increased ROS levels as measured by DHE staining (Figure 5B,D), similar to that observed after disruption of centrosome function. Together, these data are consistent with the possibility that ROS production increases in acentrosomal cells as a direct or indirect result of subsequent mitotic errors.

**Figure 5.**
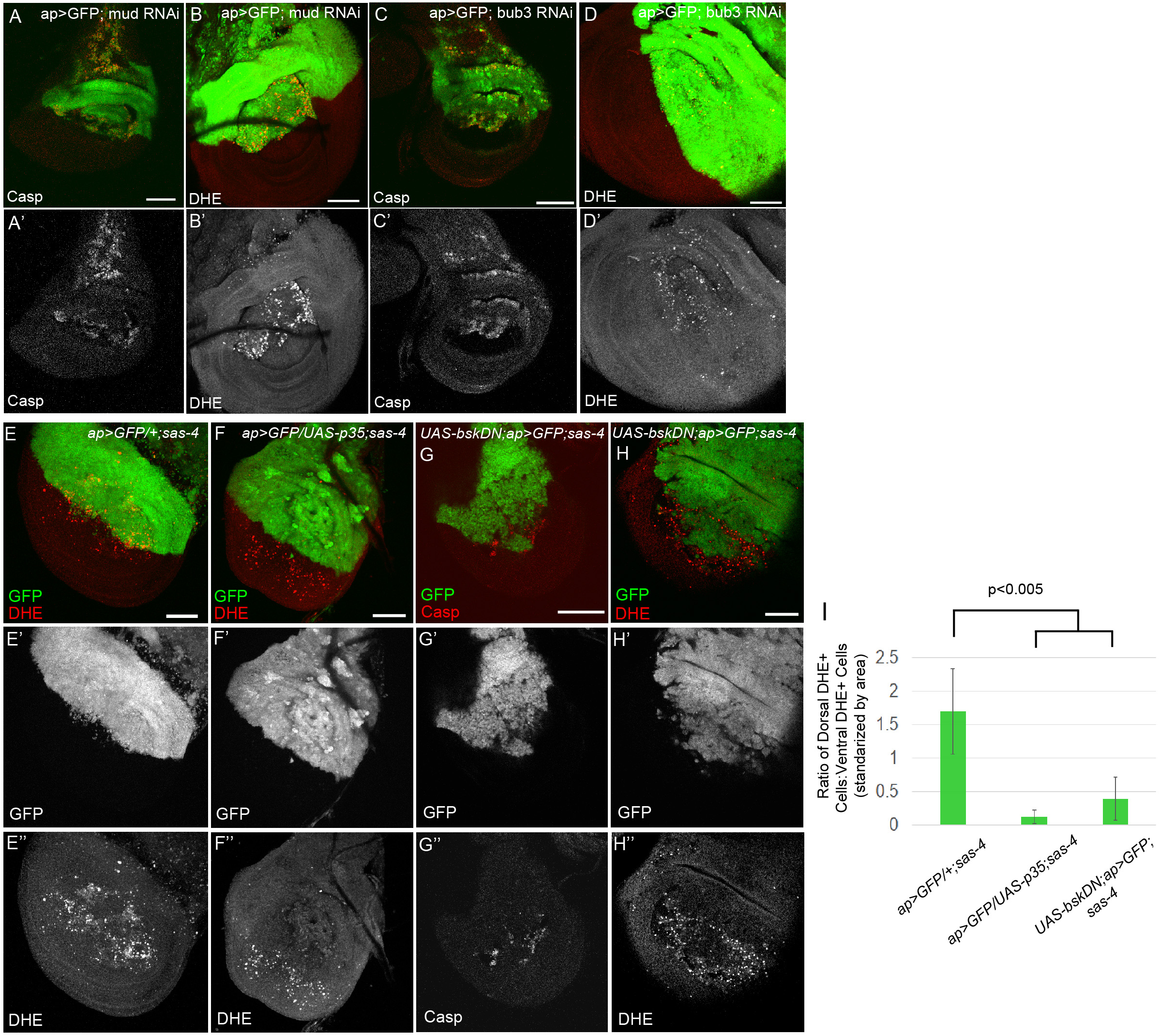
Mitotic errors induced by other stimuli also elevate apoptosis and ROS levels, and blockade of either apoptosis or JNK reduces ROS production in acentrosomal cells. (A,B) Knockdown of the mitotic spindle anchoring protein, Mud, leads to both increased apoptosis (A) and ROS (B). (C,D) Knockdown of the mitotic fidelity factor, Bub3, also increases both apoptosis (C) and ROS (D) levels. (E) *sas*-*4* homozygous mutant wing discs have elevated ROS levels, with slightly more DHE+ cells present in the dorsal compartment than the ventral (quantified in I); dorsal compartment marked by ap>GFP expression. This genotype also serves as a control for the subsequent experiments. (F) ROS production associated with centrosome loss (entire disc is *sas*-*4* mutant) is reduced by inhibiting apoptosis via p35 misexpression in the GFP+ dorsal area. (G) Misexpressing the JNK signaling inhibitor BskDN (a dominant negative form of JNK) prevents apoptosis caused by centrosome loss. (H) JNK blockade also reduces ROS levels in *sas*-*4* mutant discs. (I) Quantification of ROS levels in relevant genetic backgrounds (+/- s.d.). Blocking apoptosis through misexpression of p35 or bskDN in the dorsal region of *sas*-*4* mutant discs reduces ROS levels, relative to the ventral portion, which is *sas*-*4* mutant but does not express the indicated transgene. Scale bars are 50μm.

Recent studies in the wing disc suggest that apoptosis can induce ROS production, though this effect may be indirect (see Discussion)(SANTABARBARA-RUIZ *et al*. 2015; CLEMENTE-RUIZ *et al*. 2016; FOGARTY *et al*. 2016). To test the hypothesis that ROS induction we observed in centrosome deficient discs is a result of the apoptosis triggered by mitotic errors, we blocked apoptosis in *sas*-*4* homozygous mutant wing discs using the caspase inhibitor p35, and measured ROS levels. Blocking apoptosis significantly reduced ROS levels in *sas*-*4* mutant cells (Figure 5E,F,I). p35 blocks apoptosis by inhibiting the activity of the downstream caspase DrICE. However, it does not block activity of the upstream caspase Dronc, and this promotes continuous JNK activity in the resulting undead cells (KONDO *et al*. 2006; MARTIN *et al*. 2009). Thus the ability of p35 to block ROS elevation in *sas*-*4* mutant cells suggests that JNK activation alone is not <u>sufficient</u> to elevate ROS. (PEREZ *et al*. 2017)

The relationship between JNK signaling, apoptosis and ROS is complex. Activating JNK signaling can activate antioxidant pathways, and can also induce cell death (WANG *et al*. 2003; DHANASEKARAN AND REDDY 2017). We thus tested an alternate hypothesis: JNK signaling, while not <u>sufficient</u>, is <u>necessary</u> for the elevation of ROS we observe in centrosome deficient wing discs. To test this, we examined *sas*-*4* homozygous mutant wing discs in which JNK signaling was blocked through misexpression of bskDN. As we have shown previously, JNK inhibition via BSK-DN expression suppresses virtually all the apoptosis normally caused by centrosome loss (Figure 5G)(POULTON *et al*. 2014). DHE staining in these discs revealed a significant reduction in ROS levels after JNK blockade (Figure 5H,I), consistent with the idea that JNK signaling is necessary for elevated ROS production in centrosome-deficient wing discs. Together, these data suggest that the increased ROS levels in *sas*-*4* mutant cells depend on the completion of apoptosis, regardless of whether JNK is hyperactivated (as in the p35+ cells) or blocked (as in the bskDN cells). These results are considered further in the Discussion.

### G6PD expression buffers ROS production and G6PD knockdown elevates apoptosis caused by centrosome loss

Our earlier work revealed that mitotic errors induced by the absence of centrosomes trigger apoptosis in the aneuploid cells (POULTON *et al*. 2014). However, these data also revealed that many cells that are challenged with centrosome loss evade death, though their cell cycle is lengthened. One possibility is that some of genes upregulated in *sas*-*4* mutant wing cells help centrosome-deficient cells survive in the presence of mitotic stress. To test this hypothesis, we performed a small candidate RNAi screen testing for genetic interactions with *sas*-*4*, predicting that knockdown of genes encoding proteins that helped cells cope with centrosome loss would enhance the *sas*-*4* knockdown phenotype. The candidate genes were chosen based on their roles as key players in redox balance or related signaling pathways (Table 5). From this screen, we identified a significant interaction between *sas*-*4* and *Glucose*-*6*-*phosphate dehydrogenase (G6PD)*. While knockdown of either gene alone had minimal effects on adult wing blade morphology, knocking down both genes led to a significant increase in wing blistering (Figure 6A,B)—the *g6pd* RNAi line was previously shown to significantly reduce G6PD levels and activity (TEESALU *et al*. 2017). G6PD is the rate-limiting enzyme in the pentose phosphate pathway, converting Glucose-6-phosphate to 6-phosphoglucono-s-lactone. This reaction also generates NADPH, which is used by Glutathione reductase to produce reduced Glutathione (GSH), a potent antioxidant. Thus, in many cell types, G6PD can be a central player in the ability to limit ROS levels (STANTON 2012).

**Figure 6.**
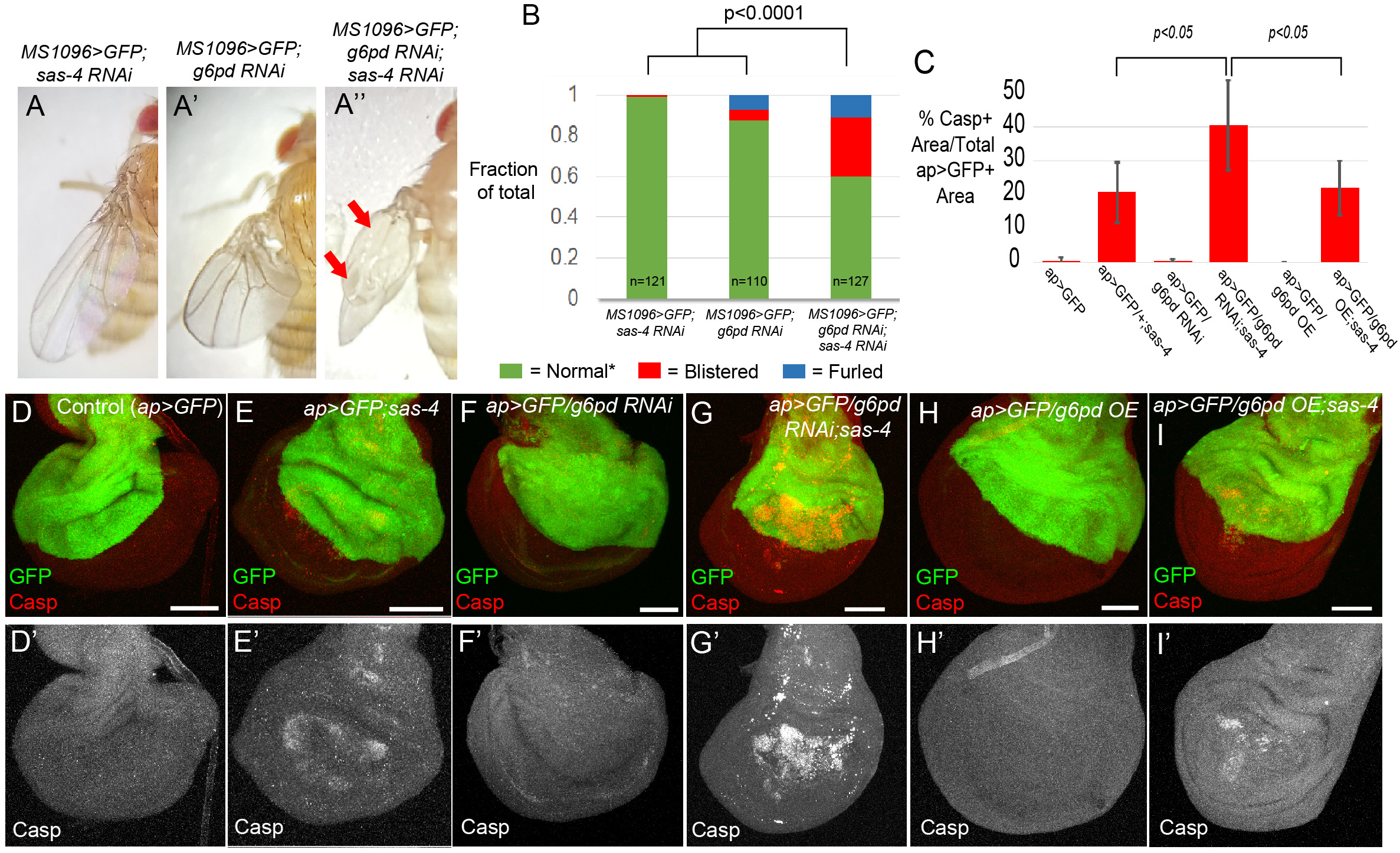
G6PD helps prevent apoptosis in acentrosomal cells. (A-C) Representative adult wing from the indicated genotypes. (A) Knockdown of *sas*-*4* with MS1096-Gal4 has no observable effect on adult wing morphology. (A’) *g6pd* knockdown has only minor effects on wing blade morphology, although the wings of these flies are held erect relative to the body. (A”) Simultaneous knockdown of both *sas*-*4* and *g6pd* leads to significantly more wing blisters and furled wing phenotypes. (B) Quantification of effects on wing blade morphology. ^*^ The “Normal” wing category includes both the WT appearance seen in the *sas*-*4* RNAi wings, as well as the erect wings with normal wing blades observed in *g6pd* RNAi flies. “Furled” refers to wings that never expanded (unfurled) after eclosion. (C) Quantification of the effects of *sas*-*4* and *g6pd* manipulations on apoptosis levels. (D-I) Assessment of apoptosis levels in 3^rd^ instar wing discs of indicated genotypes. (D) Control *ap*-*Gal4 UAS*-*GFP* (*ap*>*GFP*) wing discs have minimal apoptosis. (E) The elevated apoptosis throughout the wing pouch characteristic of *sas*-*4* homozygous mutants is not altered by expression of GFP in the dorsal region via ap>GFP. (F) Expression of *g6pd* RNAi using ap-Gal4 does not increase apoptosis in a WT background. (G) Expression of *g6pd* RNAi using ap-Gal4 in the *sas*-*4* mutant background significantly increases the incidence of apoptosis associated with centrosome loss, in the cells where *g6pd* is knocked down. (H) G6PD overexpression (OE) does not increase apoptosis in a WT background. (I) G6PD OE is not sufficient to reduce apoptosis caused by centrosome loss. Scale bars=50μm. All images are maximum intensity projections.

In our previous analysis of other *sas*-*4* genetic interactions (POULTON *et al*. 2014), an increased adult wing blistering phenotype correlated with increased levels of apoptosis during larval stages. We therefore examined apoptosis levels, using activated Caspase staining, to determine if there is any enhancement or suppression of the *sas*-*4* apoptotic phenotype after knockdown of G6PD. While *g6pd* RNAi alone led to no detectable increase in apoptosis (Figure 6D vs F, quantified in C), apoptosis was significantly increased when *g6pd* was knocked down in acentrosomal cells relative to that observed in acentrosomal cells alone (Figure 6E vs G, quantified in C). This was consistent with the hypothesis that G6PD buffers the elevated ROS production induced by centrosome loss. To directly test this, we stained for DHE in *g6pd* knockdown cells in the *sas*-*4* mutant background. Consistent with the hypothesis, we observed an even greater increase in ROS levels in acentrosomal cells that also lack G6PD (Figure 7B vs D, quantified in G), while G6PD knockdown alone did not elevate ROS (Figure 7C,G). These data suggest that elevated G6PD expression in centrosome-deficient cells helps prevent the death of some of the cells attempting to cope with the loss of centrosomes, presumably by limiting the amount of ROS.

**Figure 7.**
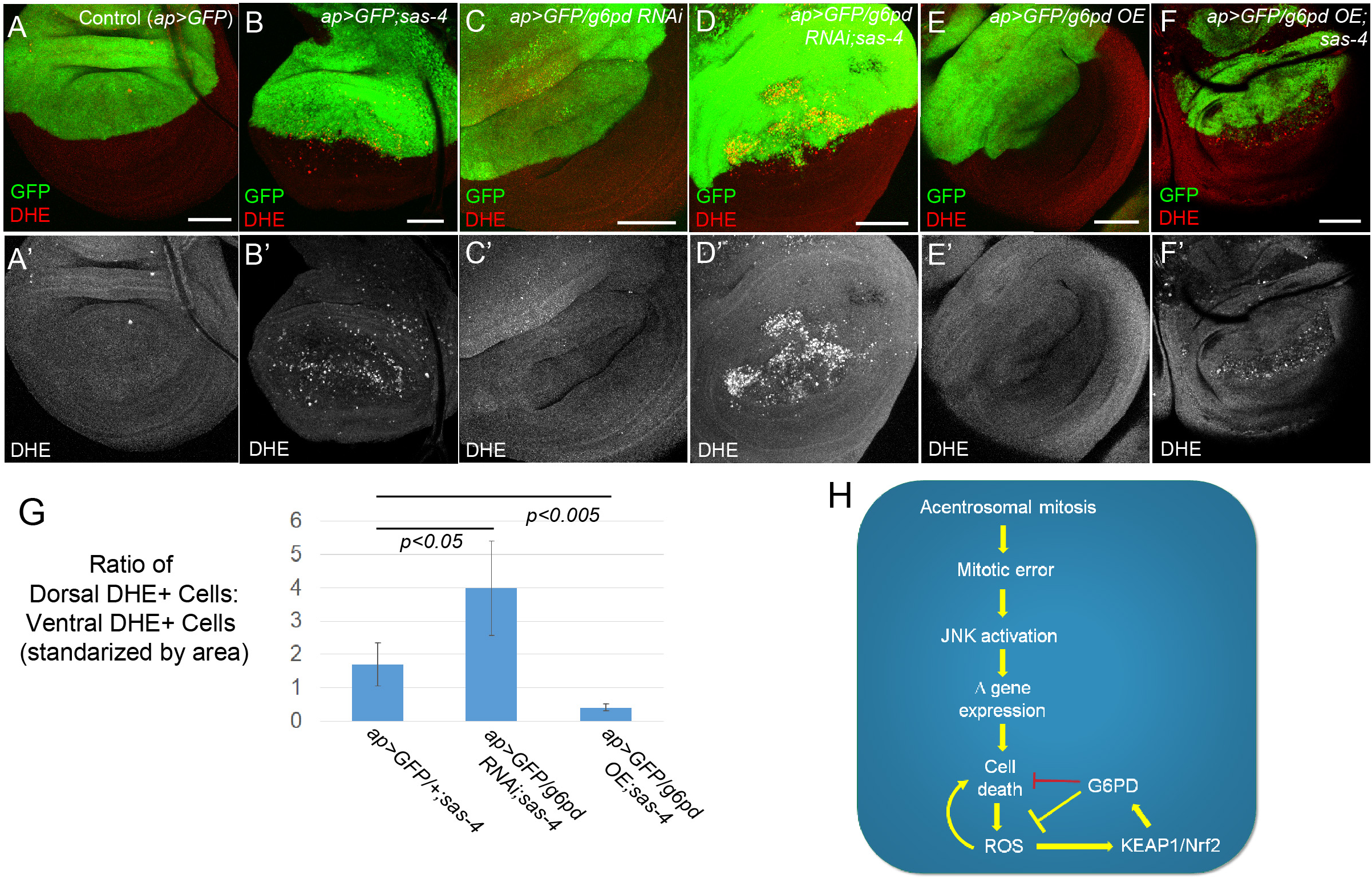
G6PD buffers acentrosomal cells against ROS production. (A-F) Assessment of ROS levels in 3^rd^ instar wing using DHE. (A) *ap*>*GFP* alone does not induce ROS. (B) *ap*>*GFP* also does not affect levels of ROS associated with centrosome loss (*sas*-*4* homozygous mutant disc). (C) *g6pd* knockdown in otherwise normal cells does not induce ROS production. (D) However, *g6pd* knockdown does elevate ROS in *sas*-*4* mutant cells above levels caused by centrosome loss alone. (E) G6PD overexpression (OE) does not affect baseline ROS production in WT. (F) G6PD OE significantly reduces levels of ROS in acentrosomal cells. Scale bars=50μm. All images are maximum intensity projections. (G) Quantification of the effects of *sas*-*4* and *g6pd* manipulations on ROS levels—see the Methods for a detailed description of these calculations. (H) Model of the relationships between the relevant pathways, processes, and genes. Because G6PD buffers ROS levels in some cells, and because ROS can contribute to cell death, G6PD can indirectly inhibit cell death [this indirect relationship is indicated by the red repression symbol].

These observations indicated that the increased expression of G6PD we observed in *sas*-*4* mutant cells, as revealed by our RNA-Seq data, might serve as a feedback response to limit apoptosis and ROS production. To test this, we overexpressed G6PD in *sas*-*4* mutants. Consistent with the role of G6PD in antioxidant generation, ROS levels were significantly reduced in the *sas*-*4* mutant cells overexpressing G6PD (Figure 7F vs B,G). Interestingly, overexpression of G6PD did not detectably affect the levels of apoptosis caused by centrosome loss (Figure 6E vs I, quantified in C). Together, these data indicate that the upregulation of G6PD is important in limiting ROS production, and reveal that although its basal level of expression helps prevent apoptosis in mitotically stressed acentrosomal cells that have not already entered the apoptotic path, increasing G6PD levels alone is not sufficient to eliminate apoptosis caused by centrosome loss (Figure 6I,C). This interpretation fits well with our observations above indicating that increased ROS levels are largely downstream of apoptosis, thus decreasing ROS in cells already firmly committed to the path to apoptosis would not necessarily be expected to decrease apoptosis in this context.

## Discussion

Transcriptional responses to cellular and tissue injury are major determinants of cell behavior and homeostasis. Drosophila imaginal discs have served as powerful models to identify the primary signaling pathways and biological processes governing these responses, and to dissect their relationships to one another (BEIRA AND PARO 2016). JNK signaling is now well-established as a central player in these events. JNK serves multiple roles, including sensing the initiating cell stress, activating pathways that can alleviate cellular stress (e.g. DNA repair pathways)(HAYAKAWA *et al*. 2004; PICCO AND PAGES 2013), triggering apoptosis when damage is severe (IGAKI 2009), and promoting activation of secondary, mitogenic signaling pathways through upregulation of their ligands, which ultimately drives large-scale processes of tissue repair such as compensatory proliferation (RYOO *et al*. 2004). More recently, in the context of cell/tissue damage, JNK has also been implicated in regulating redox balance (CLEMENTE-RUIZ *et al*. 2016; FOGARTY *et al*. 2016; KHAN *et al*. 2017), which appears to be an important aspect of the cellular and tissue-level response to that damage.

We are interested in the response to centrosome loss, and in the pathways that buffer and compensate for the mitotic defects and resultant apoptosis that centrosome loss causes in some tissues (POULTON *et al*. 2014; POULTON *et al*. 2017). To extend this analysis, we examined the transcriptional response to centrosome loss. This revealed that mitotic errors induced by centrosome loss trigger a complex transcriptional response in wing imaginal discs, including JNK-dependent changes in gene expression. Interestingly, a previous microarray-based study of transcriptome profiles in acentrosomal fly cells did not detect major changes in gene expression of multiple components of particular cellular processes (e.g., no upregulation of JNK or redox pathways)(BAUMBACH *et al*. 2012). We believe the most likely reason for this difference was that that study pooled RNA from larval wing discs and brains. As we and others have shown, larval fly brains are quite robust to centrosome loss and therefore do not noticeably activate cell stress responses, like JNK signaling (BASTO *et al*. 2006; POULTON *et al*. 2017). Thus, it is likely that the changes in gene expression occurring in acentrosomal wing discs were diluted out by the inclusion of RNA from the brains in those experiments. It is also possible that technological differences between RNA-Seq and microarray platforms also contributed to our ability to detect expression changes in numerous genes.

In our RNA-Seq analysis, multiple regulators of redox balance were significantly upregulated. This finding spurred us to investigate the oxidative stress levels of acentrosomal cells, revealing that a significant fraction of these cells have high levels of ROS. We went on to identify the upregulation of G6PD as an important component of the ability of acentrosomal cells to buffer themselves against oxidative stress. Together, our data demonstrate that error-prone, acentrosomal mitosis activates JNK signaling, leading to both induction of apoptosis and to increased ROS production (Figure 7H). Our data also revealed an important mechanism to deal with this threat—transcriptional changes in redox regulators, including G6PD, which then can feedback into the process, limiting the extent of both ROS production and cell death, and thus potentially giving cells more time to fix mitotic errors without losing the affected cells.

### ROS, JNK, apoptosis, and proliferation/repair: a complex network

Recent studies have begun to reveal the complexities of the signaling network linking ROS, JNK, apoptosis, and proliferation/repair, casting doubt on the idea that they form a simple linear pathway, and instead suggesting that the responses may vary depending on the tissue and damaging agent. It is well-established that high levels of JNK signaling can induce apoptosis in imaginal discs (IGAKI 2009). In imaginal discs, ROS production increases rapidly following cell stresses such as aneuploidy (CLEMENTE-RUIZ *et al*. 2016), and is also elevated following the initiation of the apoptotic pathway, whether it be triggered by genetic manipulations (i.e. misexpression of proapoptotic Hid or Rpr (SANTABARBARA-RUIZ *et al*. 2015; FOGARTY *et al*. 2016; BROCK *et al*. 2017; KHAN *et al*. 2017), in tumor-forming genetic models (OHSAWA *et al*. 2012; PEREZ *et al*. 2017), or by physical tissue damage (SANTABARBARA-RUIZ *et al*. 2015). However, trying to fit all the data into a single, simple linear pathway is difficult. Directly inducing apoptosis or triggering caspase activity without death can lead to ROS elevation (HUU *et al*. 2015; SANTABARBARA-RUIZ *et al*. 2015; FOGARTY *et al*. 2016; KHAN *et al*. 2017; PEREZ *et al*. 2017) but it is also well-established that elevating ROS can trigger apoptosis (CAMHI *et al*. 1995; MARTINDALE AND HOLBROOK 2002; REDZA-DUTORDOIR AND AVERILL-BATES 2016), and reducing ROS can limit the apoptotic response (O’KEEFE *et al*. 2015; CLEMENTE-RUIZ *et al*. 2016; FOGARTY *et al*. 2016). There is evidence that ROS production is induced by JNK-signaling (KHAN *et al*. 2017; PEREZ *et al*. 2017), but also evidence that ROS plays a role in activating JNK (WANG *et al*. 2003; OHSAWA *et al*. 2012; SANTABARBARA-RUIZ *et al*. 2015; CLEMENTE-RUIZ *et al*. 2016; FOGARTY *et al*. 2016; KHAN *et al*. 2017; PEREZ *et al*. 2017), and that JNK signaling can induce an antioxidant response (WANG *et al*. 2003). Finally, in some situations elevating ROS appears to reduce JNK signaling (BROCK *et al*. 2017).

The studies most relevant to our work are from the Milan lab, who examined the consequences of aneuploidy induced by disrupting the spindle assembly checkpoint or spindle assembly in eye and wing disc epithelia. The parallels with centrosome loss are striking. As with centrosome loss (POULTON *et al*. 2014), disrupting spindle assembly by reducing levels of mitotic regulators such as Rod or Bub3 elevated incidence of mitotic defects like lagging chromosomes and elevated DNA damage, and the resulting aneuploid cells are removed by apoptosis (DEKANTY *et al*. 2012; CLEMENTE-RUIZ *et al*. 2016). However, if apoptosis is blocked, highly aneuploid cells accumulate and a JNK-dependent transcriptional response is triggered. The “undead cells” elevate expression of the morphogen Wingless and drive tissue overgrowth, paralleling the effects of centrosome loss (POULTON *et al*. 2014). In a follow-up study, two more parallels were identified (CLEMENTE-RUIZ *et al*. 2016). First, dying cells accumulate elevated levels of ROS, as measured by GSTD1-GFP. Second, they also induce a transcriptional response that includes a series of JNK target genes and a set of genes involved in buffering ROS.

In other cases, the apoptosis-JNK-ROS connections are even more complex. For example, in eye imaginal discs, when apoptosis is induced by expression of Hid but death is blocked by p35 expression, the production and release of <u>extracellular</u> ROS is triggered (FOGARTY *et al*. 2016). This leads to recruitment of hemocytes that secrete the TNFα relative Eiger, which in turn activates lower and thus non-apoptotic levels of JNK activation in neighboring cells (FOGARTY *et al*. 2016). This promotes expression of mitogenic signals that contribute to processes such as compensatory proliferation/regeneration and apoptosis-induced proliferation. The generation of extracellular ROS appears to rely on plasma membrane targeted ROS generators such as Dual Oxidase (DUOX)(FOGARTY *et al*. 2016; KHAN *et al*. 2017). Intriguingly, the DUOX maturation factor NIP, encoded by *moladietz (mol)*, is itself upregulated by JNK activity (KHAN *et al*. 2017). JNK-induced extracellular ROS (produced by the NIP-DUOX mechanism) then helps maintain JNK activity in neighboring cells to drive tissue repair (KHAN *et al*. 2017).

Comparing our findings with these and other studies reveals interesting similarities and differences. First, consistent with these studies, we find that induction of apoptosis caused by loss of centrosomes or other mitotic regulators can trigger ROS production. However, our p35 data suggest that completion of apoptosis is important for ROS production in acentrosomal cells, which contrasts with data indicating that blocking apoptosis did not reduce ROS levels associated with misexpression of Hid (FOGARTY *et al*. 2016) or CIN induction (CLEMENTE-RUIZ *et al*. 2016). This difference suggests that the circuitry of these interconnected processes may be wired differently in acentrosomal cells, leading to different experimental outcomes. For example, different tissues might have different responses downstream of Dronc, which remains active after p35-mediated apoptosis blockade. Further studies will be also required to elucidate the cellular mechanisms generating ROS in these systems; NIP-DUOX is an interesting candidate (see below).

It is also worth considering what our data suggest about the relationship between JNK activity and ROS levels. As described above, recent data indicate that JNK can drive ROS production through NIP-DUOX (KHAN *et al*. 2017). However, others found that JNK is not essential for ROS production in the wing disc in response to aneuploidy (CLEMENTE-RUIZ *et al*. 2016) and that JNK activation is not sufficient for ROS elevation in the eye disc (FOGARTY *et al*. 2016). In contrast, we found that blocking JNK signaling in acentrosomal cells reduced ROS levels, which is very similar to recent findings from the Bergmann lab, who found that blocking JNK in *scrib ras^v12^* clones prevents ROS production (PEREZ *et al*. 2017). In our experiments, we speculate that the reduction in ROS levels after JNK blockade is due to the subsequent inhibition of apoptosis, because when we blocked acentrosomal cell death via p35 misexpression, which leads to hyperactive JNK signaling, we did not detect increased ROS levels, instead finding decreased ROS in the p35+ cells. Perhaps the sustained, high levels of JNK signaling in these undead cells somehow subverts the NIP-DUOX mechanism of ROS production, either directly or indirectly through massive upregulation of ROS antagonists, like G6PD. Alternately, the mechanism driving NIP-DUOX mediated ROS levels may not be active in acentrosomal cells. Indeed, despite clearly identifying the upregulation of numerous JNK targets, our RNA-Seq data did not detect any significant upregulation of *mol*, even in individual mutant gene lists (i.e., *sas*-*4* vs control or *asl* vs control). Thus it is possible that ROS production caused by centrosome loss occurs independently of NIP-DUOX. Of course, it is also plausible that differential sensitivities associated with different RNA-Seq protocols or downstream analyses may explain the absence of *mol* upregulation in our data. Thus, our data and others clearly suggest important links between cellular damage, JNK activation, initiation of the apoptotic cascade, and ROS production. However, the circuitry connecting these events and the mechanisms underlying those relationships appears to vary depending on the nature or severity of the damage. This would benefit from further exploration.

### JAK-STAT and KEAP1-Nrf2 signaling

Another signaling pathway implicated by our RNA-Seq data was the JAK-STAT pathway. In our list of upregulated genes shared by both *sas*-*4* and *asl*, we noted increased expression of Upd2 and Upd3, both ligands of the pathway and previously identified JNK targets (PASTOR-PAREJA *et al*. 2008; BUNKER *et al*. 2015; SANTABARBARA-RUIZ *et al*. 2015). This list also included inositol pentakisphosphate 2-kinase (Ipk1), which can regulate Jak-Stat signaling (SEEDS *et al*. 2015), and the JAK-STAT target gene *vir1* (DOSTERT *et al*. 2005). Indeed, we did detect enrichment of STAT92E binding sites in our set of genes upregulated in both *sas*-*4* and *asl* loss (Figure 2F), and saw upregulation of a JAK-STAT reporter in acentrosomal cells, though only when apoptosis is inhibited (Suppl Figure 1A-C). We attempted to test for genetic interactions by inducing simultaneous *upd2* and *sas*-*4* knockdown, but the severe phenotypes (massive apoptosis and abnormal adult wings) caused by knockdown of Upd2 alone precluded analysis of the interaction. Other studies demonstrate a role for JAK-STAT in regulating the compensatory proliferation and regeneration processes in damaged imaginal discs (OHSAWA *et al*. 2012; KATSUYAMA *et al*. 2015; SANTABARBARA-RUIZ *et al*. 2015; CLEMENTE-RUIZ *et al*. 2016). We speculate that the increased levels of JAK-STAT ligands we detected are likely involved in the compensatory proliferation response we previously demonstrated occurs in acentrosomal wing discs (POULTON *et al*. 2014).

We also looked for connections with the KEAP1-Nrf2 pathway, which is well known as an important regulator of cytoprotective responses to oxidative stress (SYKIOTIS AND BOHMANN 2008; LOBODA *et al*. 2016; SIES *et al*. 2017). Nrf2 is a bZIP transcription factor that positively regulates antioxidant response proteins. KEAP1 keeps it inactive by anchoring it in the cytoplasm and targeting it for proteasomal destruction, but oxidative stress relieves this inhibition (Figure 7H). G6PD is a target of KEAP1-Nrf2 signaling (LOBODA *et al*. 2016). Many of the GST family proteins that we found to be upregulated in acentrosomal cells have also previously been found to be upregulated when KEAP1-Nrf2 signaling is experimentally activated (LOBODA *et al*. 2016). Indeed, the Drosophila Nrf2 homolog, Cap-n-collar (Cnc), was recently proposed to help limit ROS levels in the wing disc following tissue damage induced by Rpr misexpression (BROCK *et al*. 2017). In their model, increased ROS levels activate Cnc, which negatively regulates ROS levels via increased transcription of ROS suppressors. They also suggest that this Cnc-mediated mechanism to restrict ROS levels helps maintain an optimal level of JNK signaling needed for tissue repair and development. Our attempts to disrupt KEAP1-Nrf2 signaling in our RNAi-based genetic interaction screen did not reveal any significant genetic interactions with *sas*-*4* knockdown (Table 5). However, whether that is due to true lack of functional interaction or a simple lack of effective knockdown remains unclear. Nevertheless, our data suggest that, G6PD upregulation serves as an important buffer against excessive ROS levels—reducing G6PD levels in acentrosomal cells significantly increases ROS levels above that induced by centrosome loss alone. Furthermore, apoptosis is elevated after loss of G6PD in acentrosomal cells, perhaps due to the increased levels of ROS. Induction of antioxidant-promoting proteins like G6PD and the GST proteins may be a part of a response by which lower levels of JNK induced in not yet apoptotic cells may allow them to survive and correct minor mitotic errors—similar to the response in neighboring cells induced by extracellular ROS (SANTABARBARA-RUIZ *et al*. 2015; FOGARTY *et al*. 2016). In contrast, the higher levels of ROS associated with loss of G6PD in acentrosomal cells may lead to higher, intolerable levels of JNK activity that pushes them down the apoptotic pathway.

### Centrosome loss, immunity, and xenobiotic detoxification

There are some remaining mysteries in the sets of genes upregulated after centrosome loss. We expected to see JNK pathway target genes, based on our earlier work, and the genes involved in the oxidative stress response and in apoptosis also seem reasonable based on the literature. However, two of the other most prominent gene sets were more puzzling: the upregulated list included many genes involved innate immunity and in in the xenobiotic response to toxic compounds (Table 3 and 6). As we considered this, we were made aware of previous work by the Ruvkun lab in C. elegans that suggests intriguing interpretations of our data. The Ruvkun group, in the course of an unrelated RNAi screen, noted a common response to knockdown of genes involved in certain core biological processes, including protein translation, proteasome function, mitochondrial function and mRNA processing (MELO AND RUVKUN 2012). Knockdown of critical components of any of these led to activation of both the innate immune and the xenobiotic detoxification responses. They and others found that similar effects were induced by pathogens or by pathogen derived or natural toxins (DUNBAR *et al*. 2012; MCEWAN *et al*. 2012; GOVINDAN *et al*. 2015). Based on these data, Melo and Ruvkun developed the intriguing hypothesis that animals evolved a conserved mechanism to detect and respond to pathogen attack on key cellular machines. Many pathogens encode small molecule or protein effectors that inhibit the function of these machines. In their hypothesis, cells thus evolved to respond to defects in the function of a key cellular machines by upregulating genes involved in the innate immune and the xenobiotic detoxification responses.

**Table 3.**
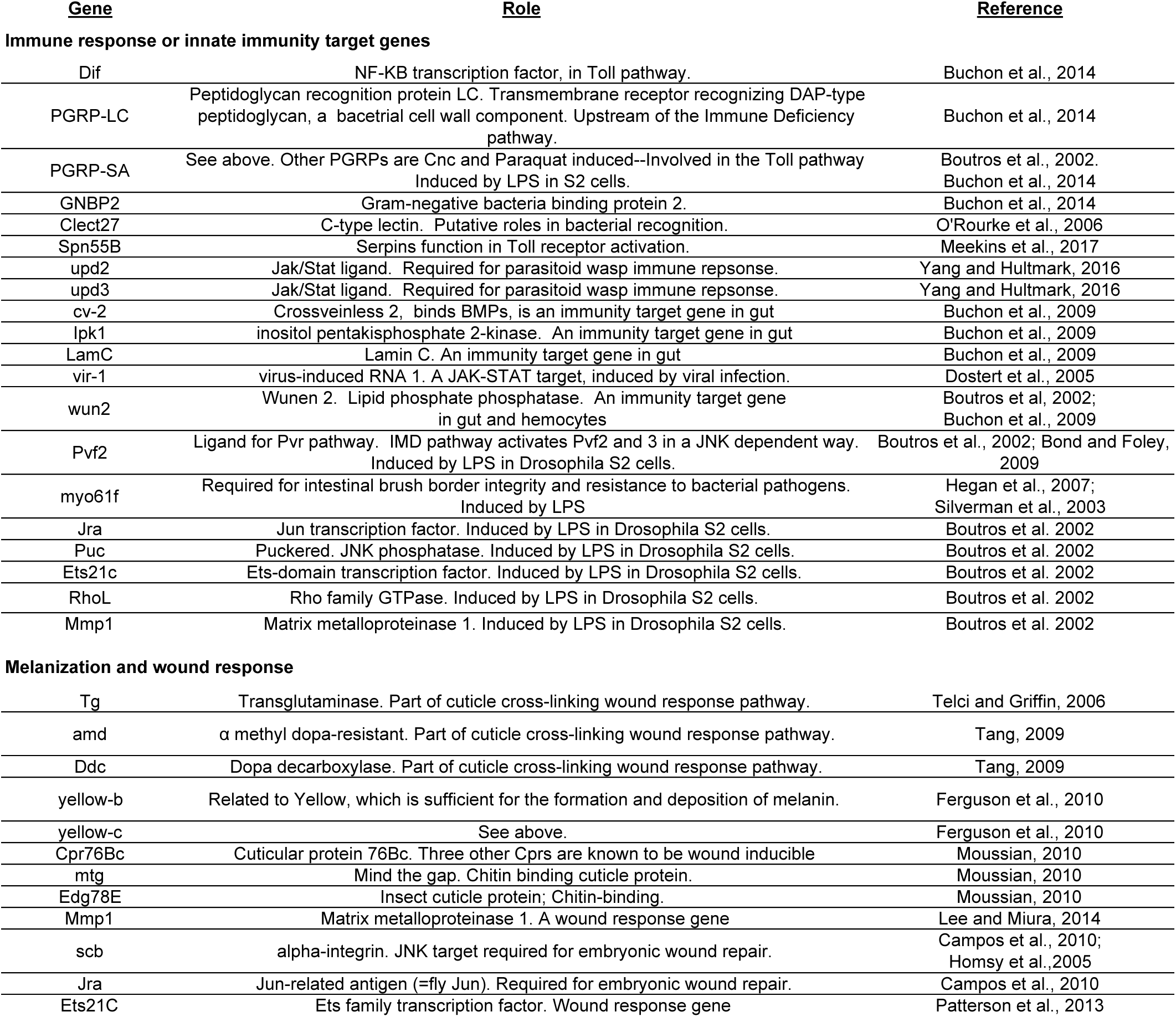
Genes with known or inferred roles in innate immunity or wound healing. Genes with reported roles in the innate immune response and/or wound healing that were significantly upregulated in both *sas*-*4* and *asl* mutants relative to WT.

**Table 4.**
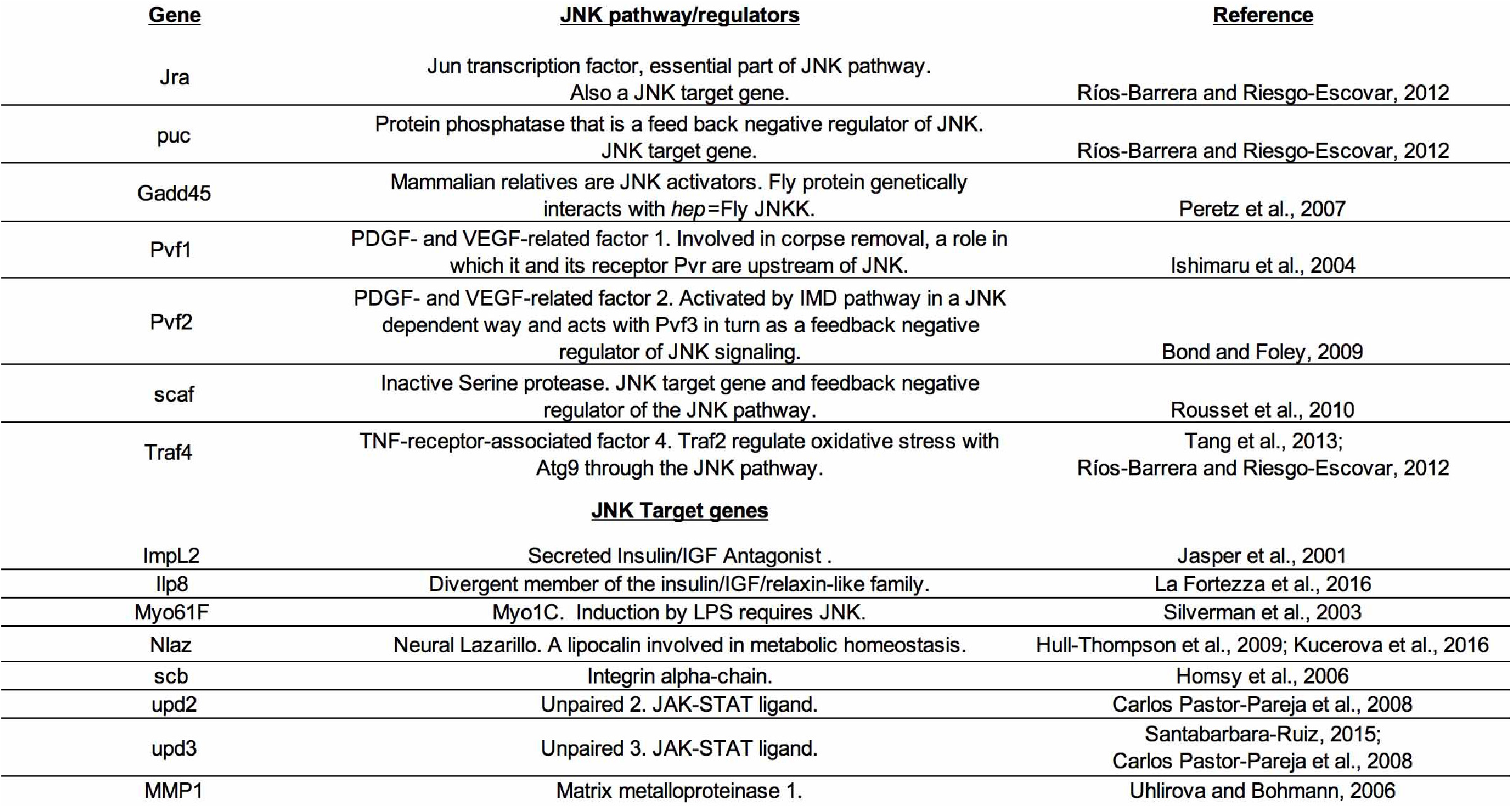
Genes related to JNK signaling. List of genes that were upregulated in both *sas*-*4* and *asl* mutants relative to WT, and are either components or the JNK signaling cascade or transcriptional targets of JNK signaling.

Our data fit well with this hypothesis. We saw upregulation of components of all three phases of the xenobiotic detoxification response, as well as many factors involved in or activated by the innate immune response (Table 3 and 6). Others have seen similar changes in response to other stresses, including the response to xenobiotic drugs like phenobarbital (MISRA *et al*. 2011) and the response to pathogenic protein aggregation (DIALYNAS *et al*. 2015; ZHAN *et al*. 2015). The genes upregulated after inactivation of the SAC also include a number encoding proteins involved in the xenobiotic response (CLEMENTE-RUIZ *et al*. 2016). Intriguing, oxidative stress also induces many of these genes (MISRA *et al*. 2011; KUCINSKI *et al*. 2017), including both ones with known antioxidant roles and those, like cytochrome p450s, without known roles in this process. Another striking parallel with the data from the Ruvkun lab is the role of the JNK pathway. They found that the responses to knockdown of genes involved in core biological processes was mediated the JNK pathway (MELO AND RUVKUN 2012), as is the case for the response to centrosome loss.

Our upregulated gene list also showed strong overlap with genes upregulated after wounding and during imaginal disc regeneration. Classic developmental biology experiments revealed that imaginal discs have a remarkable ability to regenerate after surgical or radiation damage (HAYNIE AND BRYANT 1977), by a process now known as compensatory proliferation (FAN AND BERGMANN 2008). This response also requires the JNK pathway. Strikingly, when examining the list of genes upregulated during early stages regeneration (KATSUYAMA *et al*. 2015), nine of the top twenty are also significantly upregulated after knockdown of <u>both</u> *sas*-*4* or *asl*. Thus, the genes upregulated after surgical injury and those induced by programmed cell death after centrosome loss are strikingly similar. Many of these upregulated genes are part of the generalized wound response, such as MMP1 and PGRP-SA (PATTERSON *et al*. 2013), suggesting that at least part of these transcriptional responses are parallel ones. The genes upregulated after disc regeneration also included genes involved in the xenobiotic response (e.g. three Cytochrome p450 genes and seven GST family members), suggesting the possibility that animals are genetically programmed to interpret tissue injury as a pathogen attack.

Together, our previous and current data demonstrate the robust and multi-layered mechanisms that help cells and tissues compensate for the absence of centrosomes. In proliferating epithelia, such as the wing imaginal disc, alternative microtubule nucleation pathways help build mitotic spindles, and the Spindle Assembly Checkpoint slows mitotic progression to facilitate these less-efficient pathways (POULTON *et al*. 2014). Although many cells appear to divide without obvious mitotic error, a significant number of them suffer defects related to chromosome segregation errors (e.g., lagging chromosomes, aneuploidy, and DNA damage). Our data and others suggest it is these cells that activate high levels of JNK signaling, which drives those cells into the apoptotic pathway (DEKANTY *et al*. 2012; POULTON *et al*. 2014). JNK also acts as a homeostatic buffer in the tissue by promoting compensatory proliferation in the neighboring cells (IGAKI 2009). In the present study, we uncovered an additional consequence of centrosome loss, increased ROS, and another compensatory process, upregulation of oxidative stress response genes like *g6pd*. A growing body of recent work has begun to reveal a complex network of signaling pathways and cellular processes that link cellular defects, such as mitotic errors and subsequent activation of JNK signaling and apoptosis, to oxidative stress and compensatory proliferation/regeneration. The relationships among these players is not yet fully understood, in part due to the complexity that has emerged from recent studies, including bi-directionality (e.g., ROS ←→ JNK), tissue-specific mechanisms (e.g., eye disc vs wing disc), and threshold-dependent responses (e.g., JNK promoting apoptosis vs. survival/proliferation). Future studies will help further characterize the circuitry defining these relationships and the underlying causes for the apparent specificity of some mechanisms.

## Methods

### Drosophila genetics

The following fly stocks were used: *y w* (Bloomington Drosophila Stock Center (BDSC) #1495) was used as the WT control; *sas*-*4^S2214^* (BDSC #12119), *asl^mecD^* (Blachon et al., 2008), *UAS*-*sas*-*4 RNAi* (BDSC #35049), *UAS*-*mud RNAi* (BDSC #35044), *UAS*-*bub3 RNAi* (BDSC #32989), *UAS*-*g6pd RNAi* (Vienna Drosophila Resource Center (VDRC) #101507), *ap*-*Gal4 UAS*-*GFP* (a gift from Y. Tamori, Hokkaido University), *MS1096*-*Gal4* (BDSC #8860), *en*-*Gal4 UAS*-*RFP* (BDSC #30557), *UAS bskDN* (Adachi-Yamada et al., 1999), *TRE*-*GFP* (CHATTERJEE AND BOHMANN 2012), *His2Av:eGFP* (BDSC #24163), *UAS*-*p35* (BDSC #5072), *UAS*-*g6pd[9g]* (LEGAN *et al*. 2008), *GSTD1*>*GFP* (SYKIOTIS AND BOHMANN 2008), *Ilp8:GFP^MI00727^* (BDSC #33079). A list of additional RNAi stocks tested in the candidate gene screen can be found in Table 5.

**Table 5.** Candidate genes screened for interaction with *sas*-*4*. Based on cellular function, a subset of genes was selected from the list of genes that were upregulated in both *sas*-*4* and *asl* mutants relative to WT. The stock identifiers for RNAi lines targeting the genes of interest are shown (BDSC=Bloomington Drosophila Stock Center; VDRC=Vienna Drosophila Resource Center). RNAi lines were crossed to the MS1096-Gal4 UAS-GFP (MS>GFP) wing disc driver alone and in combination with *sas*-*4* RNAi. G6PD knockdown alone had modest effects that were synergistically elevated in cells co-depleted of centrosomes by *sas*-*4* RNAi (see Figure 6A,B).

| <u>RNAi gene target</u> | <u>Stock ID</u> | <u>Crossed to *MS>GFP* alone</u> | <u>Crossed to *MS>GFP;sas-4 RNAi*</u> |
| --- | --- | --- | --- |
| Upd2 | BDSC 33949 | Strongly dysmorphic | Strongly dysmorphic |
|  | BDSC 33988 | Not determined | Normal |
| Ets21C | BDSC 39069 | Strongly dysmorphic | Strongly dysmorphic |
| Castor | VDRC 2929 | Normal | Normal |
| Cnc | VDRC 37674 | Normal | Normal |
|  | VDRC 108127 | Normal | Normal |
|  | VDRC 101235 | Strongly dysmorphic | Strongly dysmorphic |
| Dif | VDRC 30579 | Normal | Normal |
|  | VDRC 100537 | Normal | Normal |
| G6PD | VDRC 101507 | All erect, 5% blistered and 7% furled | All erect, 29% blistered and 11% furled |
|  | BDSC 50667 | Strongly dysmorphic | Strongly dysmorphic |
| ImpL3 | VDRC 110190 | Normal | Normal |
|  | VDRC 31192 | Normal | Normal |
| Jra | VDRC 31595 | Normal | Necrotic spots |
| KEAP1 | VDRC 107052 | Normal | Normal |
|  | VDRC 330323 | Normal | Normal |
| TRAF4 | VDRC 110766 | Normal | Normal |
| MMP1 | VDRC 31989 | Normal | Normal |
| WWOX | VDRC 108350 | Normal | Normal |

**Table 6.** Known or suspected xenobiotic response genes. A list of genes that have increased expression in *sas*-*4* and *asl* mutants, relative to WT, and are also proposed to function in the response to xenobiotics.

| <u>Gene</u> | <u>Role</u> | <u>Reference</u> |
| --- | --- | --- |
| **Phase 1: Cytochrome p450s** |  |  |
| Cyp18a1 | Cytochrome P450-18a1. Part of a superfamily of heme containing microsomal oxidase enzymes that metabolize both xenobiotic compounds for detoxification and endogenous substrates. | Feyereisen, 2005 |
| Cyp6a17 | Cytochrome P450-6a17. See above. | Feyereisen, 2005 |
| Cyp6a23 | Cytochrome p450-6a23. See above. | Feyereisen, 2005 |
| CG12224 | Cytochrome P450 reductase/NADPH:ferrihemoprotein oxidoreductase. Membrane-bound enzyme required for electron transfer to cytochrome P450 in the microsome from NADPH | Pandey and Fluck, 2013 |
| **Phase 2: Modify Xenobiotics to increase hydrophilicity** |  |  |
| AcCoAS | Acetyl Coenzyme A synthase. Functions in glycine conjugation of xenobiotics. Cofactor of NATs. | Montooth et al., 2006 |
| St3 | Sulfotransferase 3. | James and Ambadapadi, 2013 |
| CG15661 | UDP-glucuronosyl/UDP-glucosyltransferase involved in detoxification. | Gruenewald et al., 2009 |
| Aox (CG18522) | Aldehyde oxidase 1. In addition to metabolic roles, catalyzes the oxidation of both cytochrome P450 (CYP450) and monoamine oxidase (MAO) intermediate products. | Yang et al., 2007 |
| GSS | Glutathione synthase |  |
| GstD1 | Glutathione S-transferase D1. JNK and Nrf2 target gene. | Board and Menon, 2013; Misra et al., 2011 |
| GstD3 | Glutathione S transferase D3. Nrf2 target. | Misra et al., 2011 |
| GstE3 | Glutathione S transferase E3. Nrf2 target. | Misra et al., 2011 |
| GstE5 | Glutathione S transferase E5. | Misra et al., 2011 |
| GstE6 | Glutathione S transferase E6. Nrf2 target. | Misra et al., 2011 |
| GstE8 | Glutathione S transferase E8. Nrf2 target. | Misra et al., 2011 |
| sda | Slamdance or aminopeptidase N. Modifies glutathione-S-conjugate. gamma-glutamyl transpeptidase (GGT), aminopeptidase N (APN) and cysteine-conjugate-β-lyase (CCBL) may comprise a multi-enzyme pathway that acts on xenobiotic-glutathione conjugates. | Hausheer et al., 2011 |
| Men-b | Malic enzyme b. Induced by xenobiotics. May provide NADPH for glutathione reductase and NADPH-cytochrome c reductase. | Oda et al., 1999 |
| Zw (G6PD) | Zwischenferment. Glucose-6-phosphate dehydrogenase. A NRF2 target gene. Provides NADPH to glutathione reductase. | Loboda et al., 2016 |
| CG10365 | Glutathione-specific gamma-glutamylcyclotransferase 1. Similar to mouse Chac1. May destroy glutathione. | Bachhawat and Yadav, 2018 |
| CG18641 | Triacylglycerol lipase family. May function in metabolism of 3-phenoxybenzoic acid-containing xenobiotic triacylglycerols. | Lian et al., 2018 |
| CG4267 | Triacylglycerol lipase family. See above. | Lian et al., 2018 |
| CG7367 | Triacylglycerol lipase family. See above. | Lian et al., 2018 |
| **Phase 3: Export Xenobiotics** |  |  |
| I(2)03659 | MDR4/MOAT-B. ABC transporter-like. | Hoffman and Kroemer, 2004 |

### Detection and quantification of ROS levels

To determine ROS levels, we stained the indicated genotypes with DHE (Millipore) using the following protocol; modified from (OWUSU-ANSAH *et al*. 2008). Wandering 3^rd^ instar larvae were hemi-dissected in room temperature Schneider’s media with Pen/Strep. The inverted carcasses were immediately transferred to 1mL of Schneider’s with 30μM DHE and incubated for 10min on a nutator at room temp. The DHE solution was removed and the carcasses were washed 3x for 2min each with Schneider’s media. The wing discs were then fully dissected from the carcasses, mounted in Halocarbon oil, and immediately imaged on a Zeiss LSM Pascal confocal microscope. The same protocol was used for MitoSOX staining (5μM; Thermofisher).

To quantify and statistically compare ROS levels in the different genotypes, we used maximum intensity projections of z-stack images of DHE stainings. We then isolated and extracted the GFP+ area of the wing pouch-hinge region (i.e. the dorsal region expressing ap-Gal4 driven transgenes of interest), measured the area of the selected region, and counted the number of DHE+ cells. Because of inter-experimental variability associated with live sample preps and DHE staining, it is not possible to quantitatively compare DHE levels among different genotypes. To circumvent this issue, we used the ventral region of the wing disc as a control by which to standardize the DHE signal—the ventral compartment is homozygous *sas*-*4* mutant but does not express any of the indicated transgenes, and thus is GFP-negative. As with the dorsal region, we measured the area of the ventral wing pouch-hinge region and counted the number of DHE+ cells. We then calculated the number of DHE+ cells/area for both the dorsal (GFP+) and ventral (GFP-) regions and divided the dorsal by the ventral. Increased or decreased ROS levels induced by transgenes expression in the dorsal region will therefore alter the ratio of dorsal:ventral DHE+ cells, with each compartment separately standardized by its area. We then used Student’s t-test (Excel; two-sided, unequal variance) to determine any significant differences in means, comparing transgenic backgrounds to the *ap*>*GFP*/+;*sas*-*4* background.

### RNA-Seq experiment and analysis

Total RNA was isolated from wing imaginal discs from wandering larvae as described previously (MCKAY AND LIEB 2013). RNA from 20 larval wing discs of a given genotype was isolated. This process was repeated 3 times per genotype to yield 3 biological replicates. RNA-Sequencing was performed by the UNC High-Throughput Sequencing Facility. Libraries were created using the TruSeq Stranded mRNA kit from Illumina. Reads were aligned to the annotated dm3 *Drosophila* genome using TopHat (v2.0.14)(TRAPNELL *et al*. 2012). Read depth for each gene was generated using the Bedtools ‘coverageBed’ and ‘groupBy’ tools. Differential gene expression analysis was performed with edgeR (version 3.14.0). Differentially-expressed genes were defined as having an FDR less than or equal to 0.001, and having an average FPKM value greater than or equal to 10 in at least one sample. Gene ontology analysis was performed using DAVID (version 6.8). Motif enrichment analysis was performed using AME (MCLEAY AND BAILEY 2010), in combination with transcription factor motifs from the Fly Factor Survey database. *De novo* motif discovery was performed using DREME (BAILEY 2011). Additional details are available upon request. Graphical displays of RNA-Seq data used in figures were generated from the UCSC Genome Browser, or using edgeR’s estimates of RNA abundance.

### Immunocytochemistry and Imaging

Wing disc fixation and antibody staining were performed as previously described (ROBERTS *et al*. 2012); briefly 3^rd^ instar larvae were fixed for 20min in 4%PFA, washed 3x with 0.1%PBT, blocked with PBT plus goat serum, incubated overnight in primary antibodies, washed 3x with PBT, incubated with secondary antibodies, washed 3x with PBT, then mounted in Aquapolymount. Antibodies used: cleaved Caspase3 (1:200, Cell Signaling), MMP1 (1:50, DSHB), Jra (1:500, Santa Cruz). Alexa secondary antibodies were used at 1:500. Phalloidin was spiked into secondary antibodies at 1:500. Confocal images were acquired on a Zeiss Pascal microscope. PhotoshopCS4 (Adobe) was used to adjust levels so the range of signals spanned the entire output grayscale and to adjust brightness and contrast. Adult wing images were acquired on a Samsung Galaxy S8 attached to a Unitron FS30 microscope.

## Acknowledgements

We thank D. Bohmann, W. Orr, Y. Tamori, L. Wallrath, Bloomington Drosophila Stock Center, Vienna Drosophila Resource Center, and Developmental Studies Hybridoma Bank for reagents, J. Cuningham for technical assistance, S. Nystrom for assistance with motif analysis, T. Perdue for assistance in microscopy, the UNC High-Throughput Sequencing Facility, and our UNC colleagues for excellent suggestions. The authors declare no competing financial interests. The work was supported by NIH R35 GM118096 to MP, Research Scholar Grant 17-164-01-DDC from the American Cancer Society to DJM, and startup funds from UNC’s Department of Medicine to JSP.

## Figure Legends

**Suppl Figure 1.**
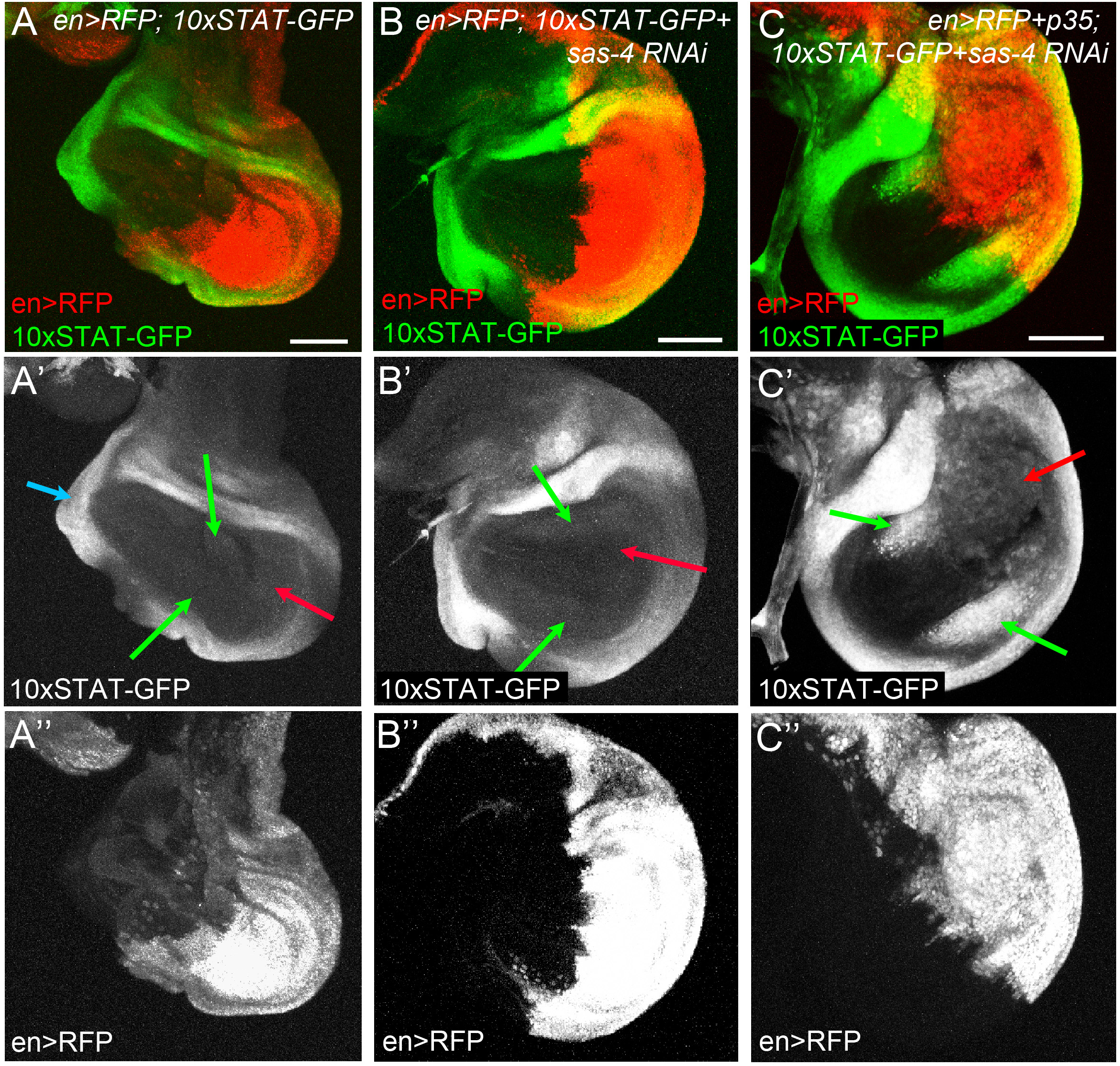
Blocking apoptosis after centrosome loss activates JAK-STAT signaling. (A) In control wing discs (en>RFP only), JAK-STAT signaling appears active only in the outer, hinge region of the disc (blue arrow), based on expression of the 10xSTAT-GFP reporter. (B) Knockdown of *sas*-*4* (i.e., centrosome loss) in the posterior half of the wing disc (marked by en>RFP+), does not noticeably affect activity of the JAK-STAT pathway—notably, there is little activity in the wing pouch (red and green arrows). (C) However, when cell death associated with *sas*-*4* knockdown is inhibited by p35 misexpression, there is a significant increase in 10xSTAT-GFP expression, particularly in the wing pouch (red arrows). Interestingly, the largest increase in JAK-STAT activity is in wildtype anterior cells adjacent to the *sas*-*4* knockdown region (green arrows in C’). Scale bars=50μm. All images are maximum intensity projections.

**Supplemental Table 1. Global gene expression data in *sas*-*4* and *asl* mutants compared to WT.**

EdgeR results comparing RNA-Seq data from *sas*-*4* or *asl* mutants to WT control wing discs; sorted by FDR. *sas*-*4* and *asl* mutant data are located in separate Worksheets.

**Supplemental Table 2. Genes up and downregulated in *sas*-*4* mutant wing discs.**

List of genes and accompanying RNA-Seq data for genes significantly (FDR<0.001) up or downregulated in *sas*-*4* mutants, relative to WT; sorted by FDR. Upregulated and downregulated genes are on separate Worksheets.

**Supplemental Table 3. Genes up and downregulated in *asl* mutant wing discs.**

List of genes and accompanying RNA-Seq data for genes significantly (FDR<0.001) up or downregulated in *asl* mutants, relative to WT; sorted by FDR. Upregulated and downregulated genes are on separate Worksheets.

**Supplemental Table 4. Genes significantly upregulated in both *sas*-*4* and *asl* mutant wing discs.**

List of shared genes significantly upregulated in *sas*-*4* and *asl* mutants, relative to WT; sorted alphabetically.

**Supplemental Table 5. Genes significantly downregulated in both *sas*-*4* and *asl* mutant wing discs.**

List of shared genes significantly downregulated in *sas*-*4* and *asl* mutants, relative to WT; sorted alphabetically.

